# Stem cell-specific ecdysone signaling regulates the development and function of a *Drosophila* sleep homeostat

**DOI:** 10.1101/2023.09.29.560022

**Authors:** Adil R Wani, Budhaditya Chowdhury, Jenny Luong, Gonzalo Morales Chaya, Krishna Patel, Jesse Isaacman-Beck, Orie Shafer, Matthew S. Kayser, Mubarak Hussain Syed

## Abstract

Complex behaviors arise from neural circuits that are assembled from diverse cell types. Sleep is a conserved and essential behavior, yet little is known regarding how the nervous system generates neuron types of the sleep-wake circuit. Here, we focus on the specification of *Drosophila* sleep-promoting neurons—long-field tangential input neurons that project to the dorsal layers of the fan-shaped body neuropil in the central complex (CX). We use lineage analysis and genetic birth dating to identify two bilateral Type II neural stem cells that generate these dorsal fan-shaped body (dFB) neurons. We show that adult dFB neurons express Ecdysone-induced protein E93, and loss of Ecdysone signaling or E93 in Type II NSCs results in the misspecification of the adult dFB neurons. Finally, we show that E93 knockdown in Type II NSCs affects adult sleep behavior. Our results provide insight into how extrinsic hormonal signaling acts on NSCs to generate neuronal diversity required for adult sleep behavior. These findings suggest that some adult sleep disorders might derive from defects in stem cell-specific temporal neurodevelopmental programs.

## Introduction

Proper brain function relies on generating diverse cell types at the appropriate time and place^1^. All neural (neurons and glia) cell types arise from a pool of progenitors called neural stem cells (NSCs)^1–8^. During development, NSCs divide to self-renew and generate distinct classes of neural subtypes over time. These processes are governed by spatial and temporal programs^9–13^. As NSCs age, they express distinct cohorts of genes; this phenomenon is called temporal patterning, allowing individual NSCs to generate distinct progeny over time^1,3–8^. The concept of temporal patterning was first defined in the *Drosophila* embryonic NSCs, which produce simple larval lineages^14^. Later, similar principles were also observed in the *Drosophila* larval NSCs that generate lineages of the adult brain^15–19^, as well as mammalian neural progenitors that generate the retina, spinal cord, and cortex^20–27^. While much is known about the temporal patterning mechanisms of the *Drosophila* NSCs, genetic programs that regulate the formation of the adult central complex (CX) lineages are poorly understood.

The insect CX is a higher-order brain center regulating complex behaviors such as navigation^28–40^, locomotion^29,41–46^, feeding^47,48^, and sleep^49–60^. The CX is a centrally located brain region comprised of four major neuropils: a handlebar-shaped protocerebral bridge (PB), a Fan-shaped body (FB), a doughnut-shaped ellipsoid body (EB), and a pair of noduli (NO)^29,36,61,62^. Two orthogonally arranged neuron types divide the CX neuropil into columns and layers: columnar (small-field) neurons divide the neuropil structure into distinct columns along the anteroposterior axis, and tangential (large-field) neurons send projections and provide input from lateral brain neuropil to the CX^36,61,62^. Recent connectome data has identified ∼400 unique neural types in the CX that are thought to be essential in regulating diverse behaviors^36^. How this diversity arises during development is not completely understood.

The neural lineages of the CX are generated by the relatively rare “Type II” NSCs^63–68^. The larval Type II NSCs occupy distinct brain regions and are organized into dorsomedial DM (1-6) and two dorsolateral (DL1-2) groups. Although only sixteen in number, Type II NSCs produce more complex and diverse lineages by generating transit-amplifying intermediate neural progenitors (INPs)^69–71^ (Figure 1A). Each Type II NSC produces roughly 40-50 INPs^65^, and each INP divides 4-5 times to produce about ten progeny; hence each Type II NSC is thought to generate approximately 400-500 progeny^63,72^. This division pattern of Type II NSCs is reminiscent of a division pattern of the primate outer radial glia (oRG) that generate lineages containing INPs and make neurons of the cortex^73–76^. Understanding how Type II NSCs produce diverse lineages of the CX might provide insights into understanding the mechanisms that regulate neural diversity in mammals. Additionally, the NSCs that generate most columnar neurons of the CX are conserved in all insects studied to date^77–84^. Thus, understanding how CX neurons are generated in *Drosophila* will likely reveal conserved developmental mechanisms.

**Figure 1.**
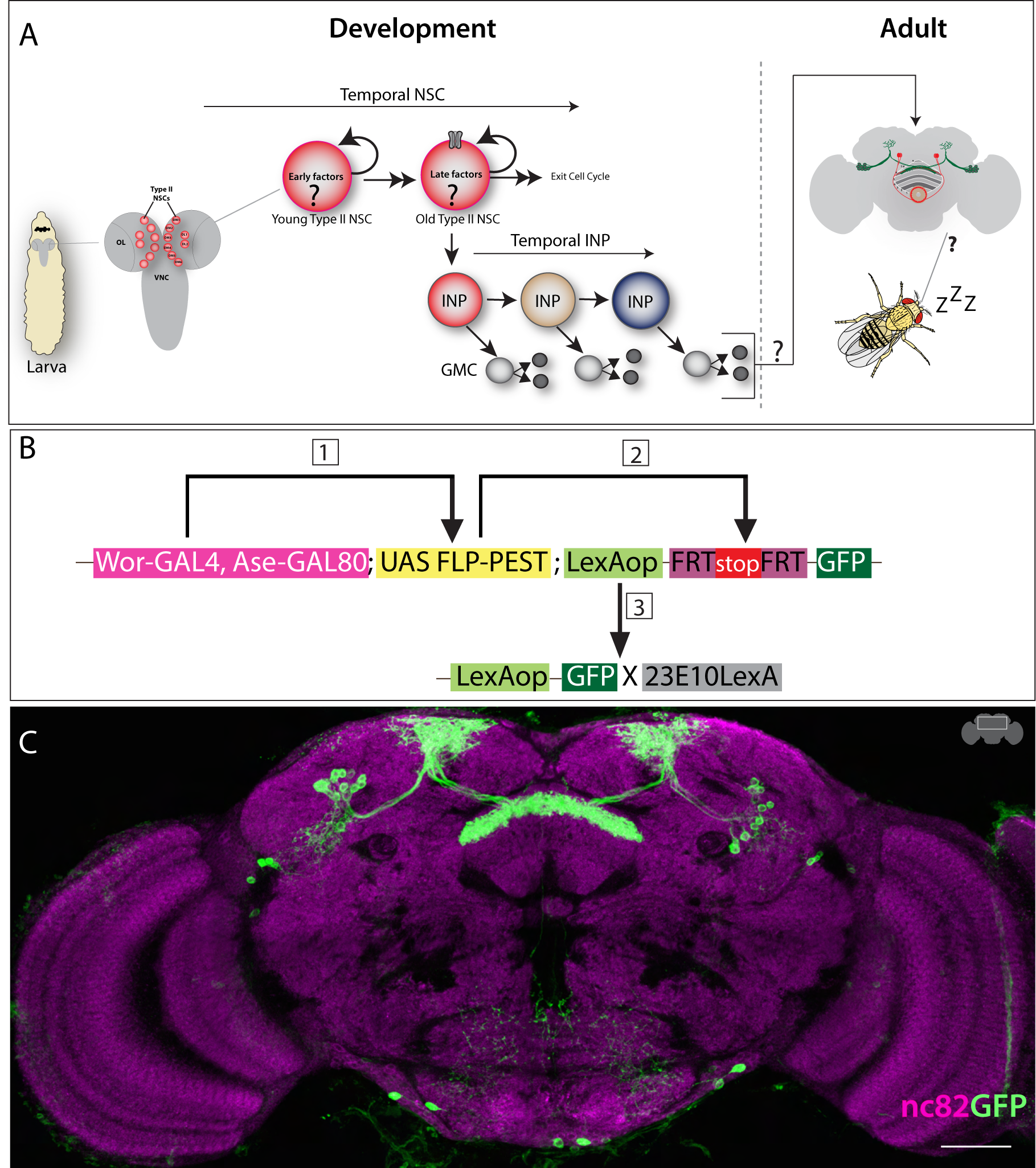
Sleep-promoting dFB neurons are generated by Type II NSC lineages. A) Schematics of larval Type II NSCs (8 per lobe: DM1-6, DL1-2), which divide asymmetrically over 120 hours ALH to generate INPs and express early and late TTFs. The temporally expressed EcR mediates the switch from early to late transition. The Type II NSC TTFs and INP temporal factors are thought to contribute to the formation and diversification of neural lineages of the *Drosophila* central complex. We are investigating the role of ecdysone signaling in the specification and function of dFB sleep-promoting neurons, which are part of the *Drosophila* sleep-wake circuit. B) Schematics showing intersectional genetic strategy for Type II NSC lineage analysis. The Worniu-GAL4, Ase-GAL80 combination drives the expression of FLP in all Type II NSCs, which excises a stop and makes LexAopmCD8GFP functional in all Type II NSCs and their progeny. This allows dFB neurons to be labeled in green if produced from Type II NSCs. C) The FLP expression in Type II NSCs labels all adult dFB neurons (green) (max projection), and nc82 labels neuropil (magenta) (projections showing only FB). The expression of GFP reporter in dFB neurons confirms that they are part of Type II NSC lineages. Scale bars, 20μm, n = 8 adult brains.

Clonal analysis has revealed unique contributions of each Type II NSC to the adult CX neuropil structures^63,64,68^. Among Type II NSCs, four (DM1-4) generate columnar neurons, and DL1 primarily generates long-field tangential neurons of the CX^63,64^. However, DM4 and DM6 also generate some long-field neurons, reflecting diverse classes of tangential input neurons^63,64^. In addition, each Type II NSCs generates distinct classes of neurons and glia over time^7,15,17,65,67,72^. Temporal clonal analysis has also revealed that INPs generate neurons of diverse identities after each division^7,65,72^, suggesting that Type II NSCs and INPs employ combinatorial temporal programs to diversify CX cell types^7,72^. What programs in Type II NSCs and INPs might regulate the temporally distinct program generating neuronal diversity? In recent studies, the larval Type II NSCs were shown to express a group of transcription factors (TFs) and RNA-binding proteins (RBPs) with precise temporal specificity^15,17^. Young Type II NSCs express early factors Castor, Sevenup, Chinmo, IGF-II mRNA-binding protein (Imp), and Lin-28; later, as NSCs age, they express ecdysone receptor (EcR), Broad, ecdysone-induced protein 93 (E93), and Syncrip^15,17,19^. Interestingly, the temporal expression of EcR around ∼55h after larval hatching (ALH) mediates early to late gene transition via NSC extrinsic ecdysone signaling^15^. Thus, unlike embryonic NSCs and larval optic lobe NSCs, generating complex adult lineages requires coordination of both stem cell intrinsic and extrinsic programs^7,15^. However, whether these temporally expressed genes and ecdysone play any role in the fate specification and function of the adult CX neurons is currently unknown. Furthermore, whether these temporal molecular cues regulate adult behaviors has remained unexplored.

Sleep is an evolutionarily conserved behavior essential for numerous physiological functions^60,85,86^. The *Drosophila* CX has repeatedly been implicated as an axis for sleep-wake regulation^50–53,57,58,87–90^. One brain area that has received particular attention is the FB of the CX, where 2310-GAL4 labeled sleep-promoting dFB neurons (subsequently called dFB neurons) innervate and regulate sleep homeostasis. These neurons are classified as long-field tangential input neurons and are about ∼12 on each side of the brain^51^. Activation of dFB neurons has been found to increase sleep^50,51^ and these neurons are more excitable following sleep deprivation^52^, although recent work has raised questions regarding the precise role of these cells^91,92^. Additionally, young adult flies exhibit increased sleep duration and depth, similar to mammals^54,55,93,94^; dFB neurons appear to play a role in generating this juvenile sleep state^54,55^. Little is known regarding the developmental origin of the dFB sleep neurons. More broadly, while linking sleep behavior to a unique NSC population is challenging in vertebrates, understanding the development of the sleep-wake circuit is crucial, as many neurodevelopmental disorders arise due to impairments in neurogenesis, circuit formation, and comorbid sleep defects with more fragmented sleep architecture^95–100^.

Here, we focus on the lineage analysis and development of the dFB neurons. Using sophisticated genetics and lineage analysis, we have identified NSCs that generate the sleep-promoting dFB neurons, known to be involved in homeostatic sleep behavior. Specifically, we show that most dFB neurons are born from the dorsolateral1 (DL1) Type II NSC, while 1-2 dFBs are born from the dorsomedial 1 (DM1) NSC. We also show that dFB neurons are generated between 48-76h ALH, and the steroid hormone signaling via E93 specifies dFB neurons. Finally, we show that NSC-specific E93 is required for normal juvenile and mature adult sleep properties. These results demonstrate the developmental origin and birth timing of sleep-promoting neurons and establish the role of steroid hormonal signaling via E93 in promoting sleep architecture.

## Results

### Sleep-promoting dFB neurons are born from Type II NSCs

Most of the neurons and glia derived from Type II NSCs populate the CX – which includes local and long-field tangential input neurons^63,64,68^. We used intersectional genetics to address whether dFB neurons originate from Type II NSCs. We used a Type II NSC-specific flippase (FLP) enzyme to permanently flip out the stop cassette from LexAop-FRT-stop-FRT-mCD8GFP and a cell class marker 23E10-LexA to label dFB neurons in the adult. In this intersectional approach, if the neurons of our interest are derived from Type II NSCs, the neurons will be labeled in green in the adult brain. We expressed FLP in all Type II NSCs using the Wor-GAL4, Ase-GAL80 driver to make the LexAop-FRT-stop-FRT-mCD8GFP reporter functional only in Type II NSCs (Figure 1B). We observed all 12 bilateral sleep-promoting dFB neurons labeled in green (Figure 1C), confirming that all sleep-promoting dFB neurons are derived from Type II NSCs. Thus, we have identified that Type II NSCs generate adult sleep-promoting neurons.

### Two distinct Type II NSCs generate sleep-promoting dFB neurons

Previous studies have assigned a unique clone morphology to each NSC present in the larval central brain, thus providing a reference framework for the clonal developmental organization of the adult *Drosophila* brain^63,64,68^. The DM1-4 Type II NSCs generate most local columnar neurons, and DL1 generates the majority of long-filed tangential input neurons with a minor contribution from DM4 and 6^63,64,101^. To determine which Type II NSCs generate dFB neurons, we used a heat shock-based lineage filtering method called cell class-lineage analysis (CLIn)^102^. This lineage filtering method classifies Type II NSC lineages by assigning cells to specific categories based on the clone morphology and connectivity patterns^102^. Using CLIn, one can generate individual Type II NSC clones by giving a temporal heat shock during development. In a successful flip-out event, the entire lineage of the Type II NSCs is labeled in red (reporter A). At the same time, the specific neuronal type marked by the GAL4 will be labeled with the GFP (reporter B) (Figure 2A, 2B).

**Figure 2.**
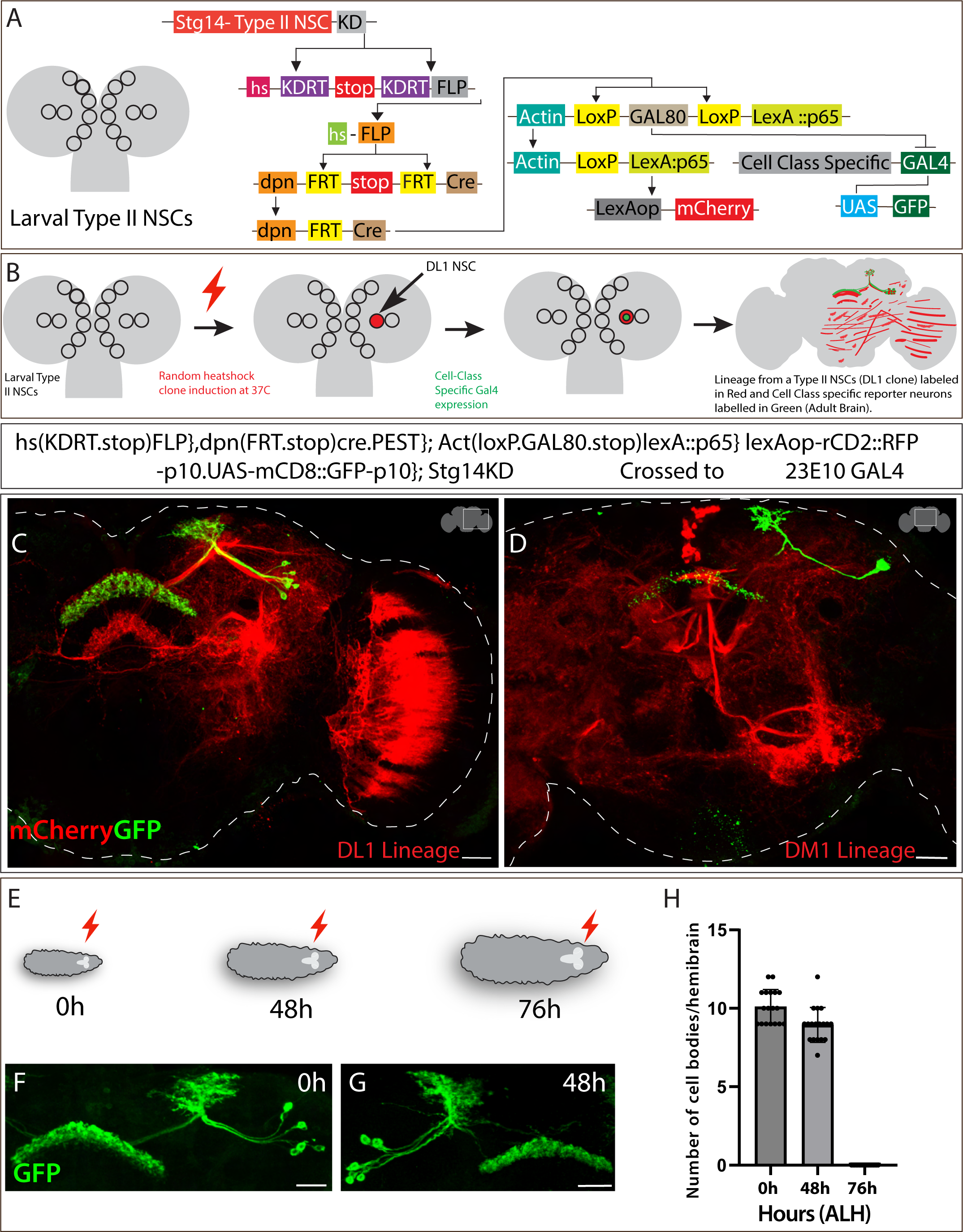
Sleep-promoting dFB neurons are generated by late DL1 and DM1 Type II NSCs. A) Schematics of CLIn intersectional genetics explaining how different genetic elements work in a sequence to label different Type II NSC lineages. The CLIn flies use Type II NSC-specific promotor, *stg,* to express KD recombinase in Type II-specific manner. The KD recombinase removes the stop sequence, bringing the FLP in frame with the heat shock promotor only in Type II NSCs. The FLP expression removes the stop sequence upon heat shock, making Cre recombinase active in Type II NSCs. The Cre recombinase makes LexA::p65 functional, making the lineage-specific expression of reporter mCherry possible. The removal of GAL80 by Cre recombinase also removes the inhibition of GAL4, making the expression of mCD8GFP in a class-specific manner. B) Schematics of how CLIn allows lineage analysis of Type II NSCs. The stochastic heat-sensitive FLP event in a single Type II NSC labels all neurons and glia born from that particular NSC. C) Single DL1 NSC clone induced at 0h ALH labels most dFB neurons (green). All the lineages from DL1 NSC are labeled in red (mCherry) and dFB neurons in green. D) Single DM1 NSC clone induced at 0h ALH labels 1-2 dFB neurons. The DM1 lineages are labeled in red (mCherry), and dFB neurons in green. E) Schematics representing heat shock given at different time points during larval development. The red lightning bolt symbol indicates heat shock given at three different time points 0h, 48h, and 76h after larval hatching (ALH). F-H) Clones induced at 0h and 48h ALH label all the dFB neurons (F, G), while clones induced at 76h ALH didn’t label any dFB neurons (not shown) H) Quantification of dFB neuron cell bodies labeled per hemibrain when clones are induced at 0h, 48h, and 76h ALH. Scale bars, 20μm, n = 16 adult hemibrains.

We adopted CLIn genetic strategy, induced a 10–12-minute heat shock at 0h ALH, and analyzed individual Type II NSC clones labeled in red (mCherry) in the adult brain (see methods for detailed protocol). Using this method, we generated single Type II NSC clones, and these individual clones were used to assign lineages to the dFB neurons. Individual clones were confirmed by comparing the reporter images with the previously published clonal map^63,64,68^. Interestingly, we observed that most dFBs (∼10) are derived from DL1 (Figure 2C), but ∼1-2 neurons are generated by DM1 Type II NSC (Figure 2D). The mixed lineage assignment of the dFB neurons to DL1 and DM1 Type II NSCs indicates heterogeneous cell types within the dFB cluster. Previous clonal studies might have missed DM1 contribution in generating long-field tangential neurons since they have not assigned these neuron types to the DM1 lineages (see discussion).

### Sleep-promoting dFB neurons are generated by late Type II NSCs

Type II NSCs express different classes of genes in the early and late portions of their lineage, which are thought to regulate the identity of the neurons born in their expression window^6,7,15,17,103^. We wanted to determine whether dFB neurons are born from early or late Type II NSCs. To identify the birth timing of the dFB neurons, we performed genetic birthdating by utilizing the heat shock-based CLIn method, as discussed above, but we performed multiple heat shock events at different times during larval development. Briefly, we crossed 23E10-GAL4, which labels dFB neurons, to the CLIn fly (for genotype, see STAR table) and gave heat shock to the progeny at three major development time points: 0h, 48h, and 76h ALH (Figure 2E). There are two phases of neurogenesis, embryonic and larval^1,6,7,104–106^. While most embryonic-born neurons die during metamorphosis^107–112^, a small population survives, undergoes metamorphosis^111,113,114^, and contributes to the adult brain^114–117^. The neurons and glia generated in the larval phase of neurogenesis predominantly contribute to adult brain circuits^63,64,68,118^. When we performed 0h ALH heat shock, we observed all the dFB neurons labeled in the adult brain, confirming that all dFB neurons are generated post-embryonically (Figure 2F). Next, we wanted to narrow down their birth time precisely to the time they are derived from Type II NSCs and performed heat shock at 48h and 76h ALH. Upon analyzing the dFB neurons in the adult brain, we observed that most dFB neurons were labeled in 0h and 48h ALH heat-shocked larvae (Figure 2F, G). However, when 76h ALH heat-shocked larvae were analyzed, we did not observe any labeled dFB neurons in the adult brain, confirming that most dFB neurons are born between 48h and 76h ALH (Figure 2E-H). Taken together, we conclude that sleep-promoting dFB neurons are born from late DL1 and DM1 Type II NSCs—at the time of EcR-mediated temporal gene expression^15^.

### Ecdysone signaling in Type II NSCs is required for dFB neuronal fate

The insect growth hormone Ecdysone regulates various stages of nervous system development, including neurogenesis, refining neural connections through pruning, and regulating programmed cell death^110,111,113,119–122^. In *Drosophila*, Ecdysone is present in varying concentrations throughout development^111,120,123^; however, Type II NSCs express EcR temporally, which transduces the extrinsic hormonal signal into the cell to regulate temporal gene expression^15^. To investigate the role of Type II NSC-specific EcR in dFB fate specification, we generated EcR-FLPStop2.0 transgenic fly. FLPStop2.0 is a modified and efficient version of the conditional loss of function strategy using the FlipStop technique^124^ (Figure 3A) with an added repeated ribozyme motif known to disrupt expression^125^ (Figure 3A, Star Methods). This method was needed due to the failure of EcR-RNAi to knock down EcR levels in Type II NSCs (data not shown). In the FLP-Stop method, tissue-specific FLP recombinase expression inverts the cassette, which results in a premature stop; as a result, a conditional loss of function allele is generated (Figure 3A). Upon this inversion, the fluorescent tag UAS*-*tdTomato becomes functional and labels mutant cells in red color^124^ (Figure 3A). To check whether EcR-FLPStop2.0 abolishes the EcR function, we expressed FLP in Type II NSCs by crossing Pointed-GAL4 to UAS-FLP in the EcR-FLPStop 2.0 background. We also added lineage tracing cassette Act-FRT-stop-FRT-GAL4 to trace mutant lineages to the adult brain (see star tables genotype). In the progeny with the genotype UAS-FLP, Act-FRT-Stop-FRT-GAL4; UAS-EcR-FLPStop2.0/Pointed-GAL4, all the EcR loss of function progeny were labeled in red, and we observed a significant reduction of EcR and E93 protein in Type II NSCs and their progeny (Figure 3S1 A-B’), confirming that EcR-FLPStop2.0 severely reduces EcR function. In the EcR-FLPStop2.0 background, we used 23E10-LexA, LexAop-mCD8GFP to label dFB neurons in green. In the progeny with the genotype UAS-FLP, Act-FRT-stop-FRT-GAL4; UAS-EcR-FLPStop2.0, 23E10-LexA; Pointed-GAL4, LexAop-mCD8GFP, we were able to simultaneously create EcR loss of function in Type II NSCs and label dFB neurons in the adult. We examined whether loss of EcR in Type II NSCs leads to defects in the specification, morphology, or connectivity of the dFB neurons. There are ∼12 dFB neurons on each side of the brain, with cell bodies located in the exterior of the CX in the PPL region. These neurons send the axonal projections to layer 6 of the dorsal FB, where they are thought to connect with the helicon cells to regulate sleep homeostasis^51^. In most experimental flies, depletion of EcR in Type II NSCs resulted in the loss of dFB neurons in the adult brain (Figure 3C-D’), suggesting that ecdysone signaling in Type II NSCs is necessary for the formation of dFB neurons.

**Figure 3.**
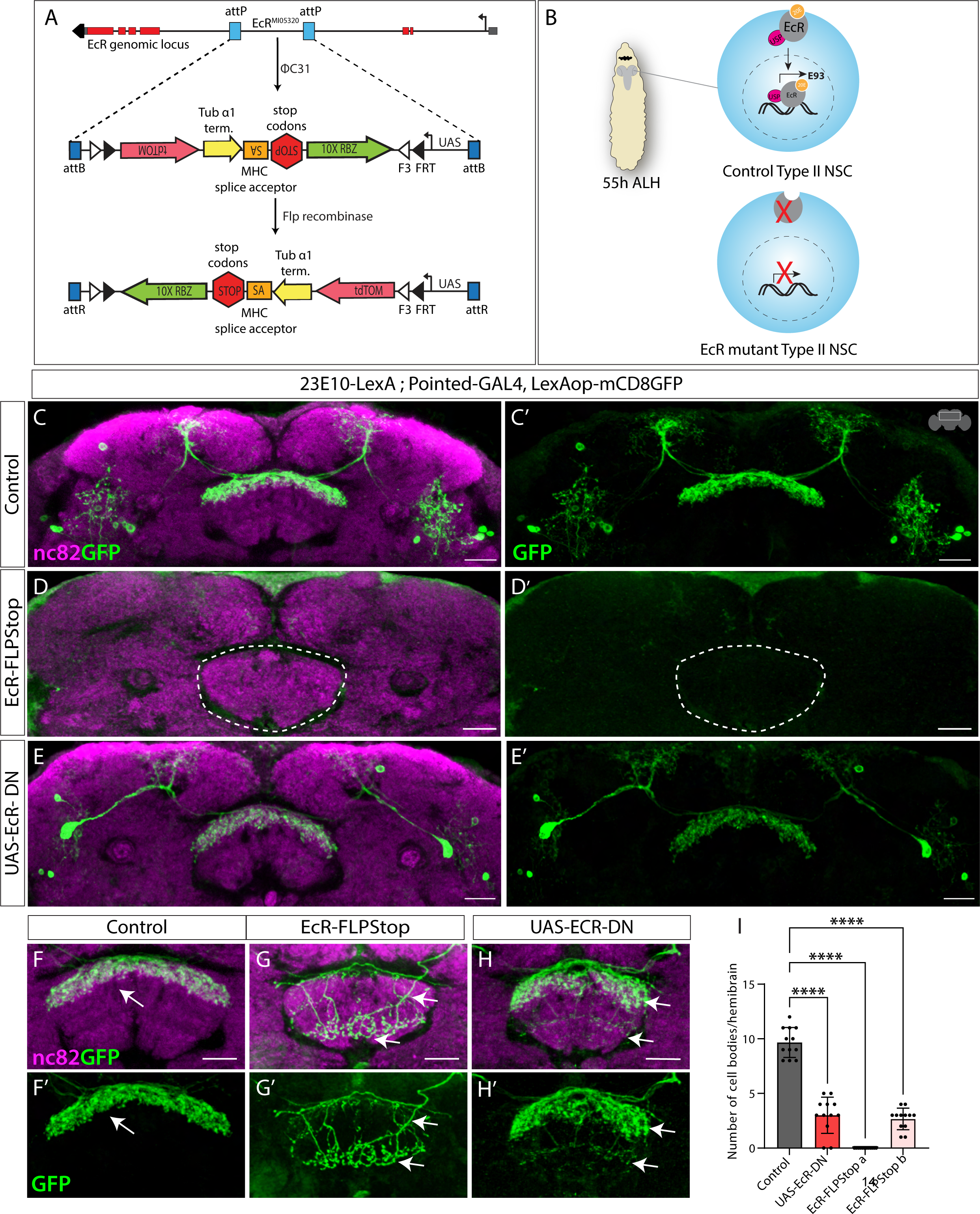
Ecdysone signaling regulates dFB neuron specification. A) Genetic elements of EcR-FlpStop2.0 scheme. The expression of FLP recombinase in Type II NSCs flips the tdTomato sequence in frame with a UAS promoter, allowing it to label mutant cells specifically under the control of the Type II specific GAL4 in red color. The FLP event also inverts the STOP sequence - transcription-based disruption (Tubα1 terminator and 10x Ribozyme sequence) and translation disruption (MHC splice acceptor paired with STOP codons) - to generate a premature stop, disrupting EcR expression and function. B) Schematics showing typical Type II NSC expressing EcR at 55h ALH that leads to expression of other EcR-induced downstream genes; upon removal of ecdysone receptor, the expression of target genes is disrupted. C, C’) Shows a control brain with GFP-labeled dFB neurons and their projection pattern in the FB. D, D’) Upon EcR loss of function in Type II NSCs, dFB neurons are not specified (dashed line annotates the FB). E, E’) Blocking ecdysone signaling in Type II NSCs using EcR-DN results in significant loss of dFB neurons. F-H’) In control brains, the dFB neurons project to layer 6 of the FB (F, F’); in EcR loss of function (G, G’) and EcR-DN (H, H’) animals, the surviving dFB neurons miss target to the ectopic FB layers indicated by arrows. I) One-way ANOVA test quantification of dFB neuron cell bodies. Error bars represent SEM; * p<0.05, **p<0.01, ***p<0.001, **** p<0.0001, NS, non-significant. Scale bars, 20μm, n = 10 adult hemibrains.

To further investigate whether the steroid hormone ecdysone regulates the dFB fate and connectivity, we use EcR dominant negative (DN)^126^ to block the ecdysone signaling. We specifically blocked ecdysone signaling in larval Type II NSCs by expressing EcR-DN with Pointed-GAL4 and assayed the fate of dFB neurons in the adult brain. We found that blocking ecdysone signaling in Type II NSCs affects dFB fate, similar to the EcR loss of function (Figure 3 E, E’). In EcR loss of function animals, we also found some animals with a less penetrant phenotype, where a few neurons were present (Figure 3F-G’); however, interestingly, the connectivity of the surviving neurons was severely disrupted (Figure 3G, G’). Compared to the control brains (Figure 3F-F’), where dFB neurons send their axonal projections to layer 6 of the dorsal portion of the fan-shaped body, in EcR loss of function (Figure 3G-G’), surviving neurons ectopically project to the ventral layers of the FB. A similar mistargeting phenotype was observed in the surviving dFB neurons in EcR dominant negative experimental animals (Figure 3H, H’), indicating multiple roles of EcR and ecdysone signaling in specifying sleep-wake circuit (see discussion). Taken together, our findings suggest that Type II NSC-specific ecdysone signaling is essential for the proper formation and identity of the dFB neurons (Figure 3I). Upon misexpression of EcR in Type II NSCs throughout development, we did not observe any increase in the dFB numbers (Figure 3S2 A-C), indicating that EcR alone might not be sufficient for generating dFB neuron types. Taken together, our studies link an extrinsic hormonal signal to the NSC-intrinsic gene programs via temporal expression of EcR in late NSCs to generate sleep-promoting neurons.

Using nc82 staining, we also investigated whether EcR in Type II NSCs regulates the overall architecture of CX neuropil structures—PB, FB, EB, and NO. Interestingly, the adult EB structure was defective upon EcR loss of function in Type II NSC lineages (Figure 3S3 A-B’). The EB did not fuse appropriately in the experimental flies, and there was a cleft in the posterior EB. We did not observe any severe morphological defects in the other neuropil structures, suggesting that ecdysone signaling regulates the fate of CX neuronal subtypes, in particular dFB neurons (Figure 3S3 A-B’). To our knowledge, these findings are the first to relate the extrinsic ecdysone signal to stem cell intrinsic gene programs to specify sleep-promoting neurons and CX development.

### Ecdysone signaling in Type II NSCs governs dFB neuronal fate via E93

Next, we wanted to understand how Type II NSC-specific ecdysone signaling regulates dFB neuronal specification. Our previous work identified ecdysone signaling as the primary regulator of early to late gene transitions in Type II NSCs^15^. Around ∼55 ALH, temporal expression of EcR in Type II NSCs activates late genes (Figure 1A), which includes the Ecdysone-induced gene, E93. Interestingly, all sleep-promoting dFB neurons express E93 in their cell bodies, indicating that EcR might specify dFB fate via E93 (Figure 4A-A’’). To test this hypothesis, we used Pointed-GAL4 to knock down E93 in all Type II NSCs during development and assayed dFB neurons labeled with 23E10-LexA, LexAop-mCD8GFP in adults. We confirmed the efficiency and specificity of UAS-E93RNAi by staining larval Type II NSCs for E93 expression (Figure 4S1). For the control experiments, we crossed Pointed-GAL4 to an empty RNAi, UAS-KKRNAi, which has the same genetic background as UAS-E93RNAi (see methods). Compared to the controls (Figure 4B-B’), reducing E93 levels in Type II NSCs resulted in the absence of all dFB neurons (Figure 4C-C’), quantified in (Figure 4E), confirming that late Type II specific E93 expression is essential for the specification of dFB neurons. In the experimental flies, we noticed 2-3 cell bodies were always present; however, we observed them in the controls as well, and these neurons are not part of the dFB cluster since they do not send projections to the dorsal fan-shaped body. Taken together, we conclude that the late temporal expression of E93, activated by EcR, regulates the formation of adult dFB neurons. To ensure the phenotype is E93 specific, we further confirmed our results using E93 RNAi without dicer and an additional independent E93 RNAi line (Figure 4S2 A-C; see STAR Methods). Both conditions produced similar phenotypes; however, E93 RNAi with dicer produced a more severe phenotype (Figure 4S2 A-C). Unlike EcR loss of function, E93 knockdown in Type II NSCs did not affect the morphology of the adult CX neuropil (including EB) (Figure 4S3 A-C’’), suggesting that E93 is not essential for the overall CX neuropil development and specifically regulates dFB neuronal fate.

**Figure 4.**
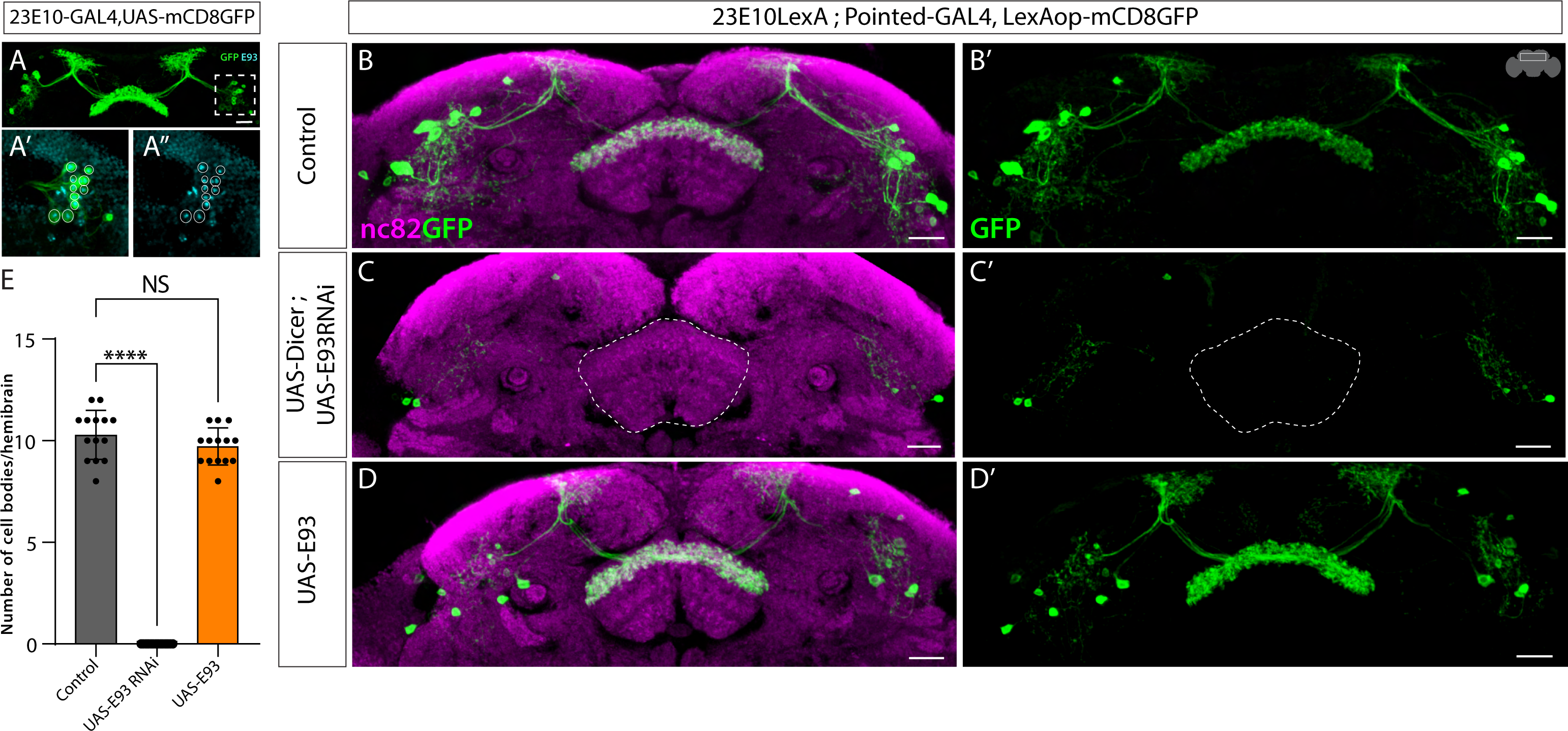
Ecdysone signaling regulates dFB neural fate specification via E93. A-A’’) The dFB neurons labeled in green (A) express E93 in cell bodies (A’, A’’). B, B’) Control dFB neurons project normally to the FB. C, C’) Complete loss of dFB neurons upon E93 knock-down in Type-II NSCs. D, D’) No change in cell body number and morphology of dFB neurons upon E93 overexpression in Type-II NSCs. E) One-way ANOVA test quantification of dFB neuron cell bodies per hemibrain. Error bars represent SEM; * p<0.05, **p<0.01, ***p<0.001, **** p<0.0001, NS, non-significant. Scale bars, 20μm, n = 14 adult hemibrains.

Next, we wanted to test whether E93 is sufficient to specify dFB neurons. We used Pointed-GAL4, UAS-E93 to miss express E93 in Type II NSCs throughout larval development and 23E10-LexA, LexAop-mCD8GFP to label dFB neurons. Compared to the control (Figure 4B-B’), we did not observe any extra dFB neurons in experimental animals (Figure 4D-D’), quantified in (Figure 4E), indicating that E93 is not sufficient to generate the dFB neuronal fate and rather a combination of factors might be involved.

Type II NSCs express EcR beginning around ∼55h ALH; E93 expression starts right after this developmental time point at a low level, which peaks around pupal formation at 120h ALH^15^. Next, we aimed to narrow down the developmental timing of the E93 function. Since dFB neurons are born between 48-76h ALH, we used the temporal and regional gene expression targeting (TARGET) system^127^ to restrict the E93 knockdown during that specific developmental time window (Figure 5A, B). Briefly, at the permissive temperature (18C), GAL80ts is active and inhibits GAL4 activity by binding to its activation domain; this inhibition is lost upon temperature shift to 29C, which allows GAL4 to function (Figure 5A). The knockdown of E93 at 48h ALH or later significantly decreased total dFB neuron number and recapitulated the phenotype observed with constitutive E93 knockdown (Figure 4C-C’), confirming that temporal expression of E93 in Type II NSCs between 48h – 76h ALH is essential for specifying dFB neurons. For the control experiment, we used UAS-E93RNAi without GAL80ts, UAS-E93RNAi; or GAL80ts constitutively grown at 29C and 18C (Figure 5C-F’). The analysis and quantification of control and experimental animals (Figure 5G) suggest that ecdysone-induced, Type II NSC specific temporal E93 expression between 48h-76h ALH specifies dFB neurons.

**Figure 5.**
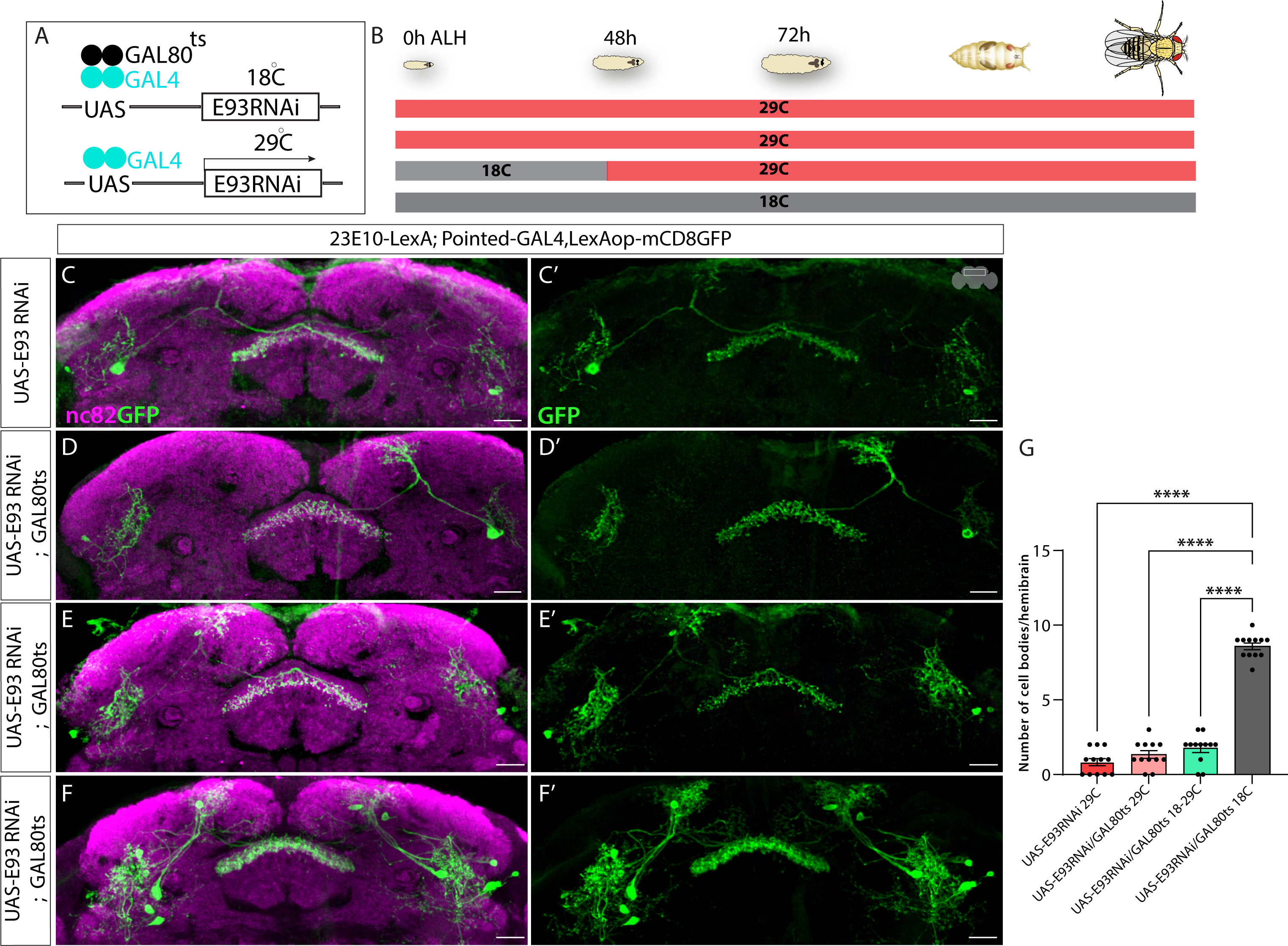
E93 expression in a restricted time window regulates dFB neuronal fate. A) Schematics of TARGET system showing GAL80ts mediated restricted knockdown of E93. At 18^0^C, GAL80ts will be active, preventing the expression of E93RNAi by inhibiting Pointed-GAL4. At higher temperatures, 29^0^C, GAL80ts will be inactive, allowing the expression of E93RNAi temporally. B) Schematics of the experimental setup showing E93 RNAi flies growing at different temperatures and under different conditions across the larval life cycle from 0h ALH to 120h ALH. The E93RNAi experimental flies containing GAL80ts were initially grown at 18^0^C and later shifted to 29^0^C around 40h ALH to make E93 RNAi expression possible in late Type II NSCs. C, C’) Shows loss of dFB neurons labeled with GFP at 29^0^C upon E93 knockdown (E93 RNAi without dicer) D, D’) Loss of dFB neurons can be seen upon E93 RNAi combined with Gal80ts grown continuously at 29^0^C (GAL80ts will be inactive at 29^0^C) E, E’) Significant loss of dFB neurons can be seen when the UAS-E93 RNAi was restricted to the late Type II NSCs using GAL80ts; the flies were grown at 18^0^C till 40h ALH and then shifted to 29^0^C to make GAL80ts inactive. F, F’) The dFB neuron number is normal when flies expressing UAS-E93RNAi combined with Gal80ts are grown continuously under 18^0^C. G) One-way ANOVA test quantification of dFB neuron cell bodies per hemibrain. Error bars represent SEM; * p<0.05, **p<0.01, ***p<0.001, **** p<0.0001, NS, non-significant. Scale bars, 20μm, n = 12 adult hemibrains.

### Loss of E93 in larval NSCs impairs adult sleep behaviors

The *Drosophila* CX has been implicated as a center for sleep regulation^36,50–53,58–60,88–90^, with evidence of a specific role for the dFB in sleep homeostasis^52,87^. We tested if loss of E93 in Type II NSCs affects sleep in adulthood. E93 knockdown in Type II NSCs using Pointed-GAL4 had a very mild effect on total sleep duration in mature adults (Figure 6A), despite dramatic effects on the dFB neuronal fate. Compared to both parental controls, there was a small reduction in daytime sleep duration and no consistent change during the night (Figure 6A). This finding is consistent with studies showing that dFB inhibition has little impact on sleep duration in mature adulthood^55,92^. We did, however, observe a substantial impact on sleep fragmentation during night, with an increase in sleep bout number (Figure 6B) and a decrease in bout length (Figure 6C) in E93 RNAi flies compared to both controls. Recent evidence indicates that sleep in *Drosophila* is not a homogenous state and that deeper sleep occurs during periods of consolidated sleep consisting of long sleep bouts^128–134^. We next asked if long bouts of sleep (60 minutes or longer) might be selectively affected by E93 knockdown; our analysis focused on the night period since this is when deep sleep primarily occurs^128,130,131^. Indeed, we found a marked reduction in the duration of sleep comprised of long sleep bouts in the night with E93 knockdown compared to controls, suggesting a role for the dFB in deep sleep (Figure 6S1).

**Figure 6.**
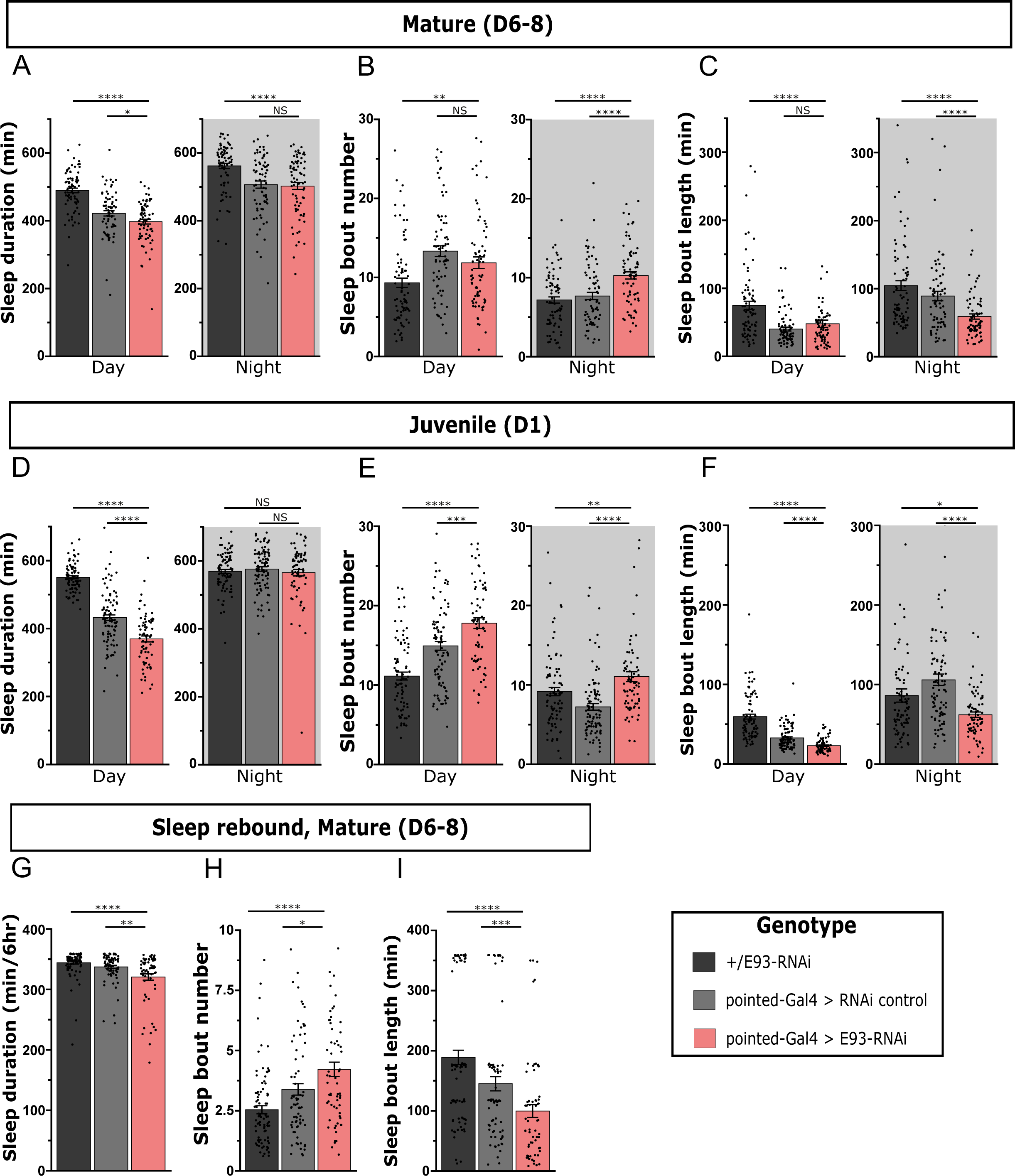
Knockdown of E93 in larval Type II NSCs impairs adult sleep. Quantification of day and night sleep duration (A), sleep bout number (B), and sleep bout length (C) in mature adult flies expressing E93-RNAi under control of pointed-GAL4 (red) compared to genetic controls (black, gray). n=79,74,74 from left to right. Quantification of sleep duration (D), sleep bout number (E), and sleep bout length (F) in juvenile adult E93-RNAi flies and controls. n=85,94,74 from left to right. Quantification of sleep duration (G), sleep bout number (H), and sleep bout length (I) in mature adult E93-RNAi flies and controls over 6 hours following a night (12 hours) of sleep deprivation. n=93,82,67 from left to right. Error bars represent SEM; *p<0.05, **p<0.01, ***p<0.001, ****p<0.0001, NS, non-significant by One-way ANOVA with Mann-Whitney multiple comparisons test corrections.

During juvenile developmental periods across species, including early adulthood in *Drosophila*, sleep depth (and duration) are elevated^54,55^. Juvenile flies (Day 1 post eclosion) exhibit increased arousal threshold during the day and night compared to mature adults (Day 5-9) and an increase in sleep duration during the day period^54,55^. We next investigated if the knockdown of E93 in Type II NSCs affects juvenile adult sleep. We found that expression of E93 RNAi was associated with a reduction in total daytime sleep duration in juvenile adults compared to controls, with no impact on sleep time during the night (Figure 6D). However, sleep was more fragmented across day and night (Figure 6E, F), as evidenced by increased sleep bout number and reduced bout length (Figure 6E,F). These results suggest a particular sensitivity of juvenile sleep to the loss of sleep-promoting dFB neurons.

Finally, we examined how knockdown of E93 in Type II NSCs affects sleep rebound following a night of sleep deprivation in mature adult flies, which is another high sleep pressure state. Total sleep duration during the rebound period (first 6 hours of the morning) was modestly reduced in Pointed-GAL4 > E93 RNAi flies (Figure 6G) but sleep during this period was more fragmented (shorter sleep bouts) (Figure 6H, I). Notably, recent work has raised the possibility that sleep relevant neurons labeled by the 23E10 driver reside in the ventral nerve cord (VNC) rather than (or in addition to) neurons in the brain^91,92^. We found that E93 knockdown in Type II NSCs impairs brain dFB neurons but spares the 23E10+VNC neurons (Figure 6S2). Our developmental approach thus assigns the brain-specific sleep roles for dFB neurons. Specifically, behavioral findings support a role for 23E10+ dFB neurons in promoting consolidated sleep in situations associated with high sleep pressure.

## Discussion

Generating complex behaviors requires the integration of various sensory modalities to generate motor output in a context-dependent manner. Distinct neuronal cell types provide unique sensory inputs or modulate motor outputs to regulate behaviors such as navigation, feeding, and sleep. Understanding brain function thus requires investigating developmental programs that establish circuits and behaviors. Here, we have investigated the lineage-specific development of sleep-promoting neurons that are an essential part of a sleep-wake circuit^51,53^. Our studies have identified the NSCs that generate dFB sleep-promoting neurons of the brain and mapped their birth timing (Figure 2C, 2D). Our work has identified the crucial role of temporal steroid hormonal signaling in regulating the specification of the dFB neurons via E93 (Figure 4). Furthermore, we identified the role of E93 in Type II NSCs in regulating juvenile and mature adult sleep (Figure 6). This study provides a new perspective on the genetic and developmental basis of sleep, and links sleep behavior to distinct NSCs.

### Lineages to sleep circuits

Diverse classes of large field input neurons feed sensory information to the CX by innervating and making connections in the different layers of the FB. Recent connectome data has specified the distinct modules in the FB layers to specific behaviors, with layers 6-7 as a sleep-wake module^36^. The FB is the most diverse structure of the CX neuropils, which receives input from the periphery via the long-field neurons that innervate different FB layers topographically. The FB long-field neurons arise from diverse NSCs, with significant contributions from the DL1 Type II NSC and DALc12v Type I NSC^63,64,68,101^. The sleep-promoting dFB neurons are long-field input neurons that innervate layers six-seven of the FB^36^. What NSC populations make the long field tangential input neurons of the sleep-wake circuit? Is the lamination of the FB birth order related? Here, we show that the dFB neurons are born from Type II NSCs, while the majority are derived from the DL1 lineages; a few are born from the DM1 lineage. Previous clonal studies could not find any contributions of DM1 lineage to the long-field input neurons. However, consistent with our findings, a recent connectomics-based lineage assignment study has also proposed DM1 lineage contribution to the FB tangential input neurons^101^. Although all ∼12 neurons look similar in morphology, our clonal analysis data suggests a likely heterogeneity of cells in this cluster. It could be each neuron in that cluster is a unique neuron type getting inputs from distinct upstream neurons and part of unique circuits. The connectomic-based studies will help in assigning unique input and output neurons to these distinct classes of dFB neurons. Furthermore, single-cell RNA-Sequencing will help delineate the distinct cell types in the dFB cluster.

Are the neuron types innervating distinct FB layers specified at different times during development? Our findings reveal that the sleep-promoting dFB neurons are born from the old Type II NSCs between 48-76h ALH. Previous studies indicate that large field tangential input neurons that innervate ventral FB layers are generated only until 72h ALH, indicating the time-dependent lamination of the FB^17,135^. Once specified, the dFB neurons innervate the FB around 48h APF where they intermingle with arousal promoting dopaminergic inputs, a process regulated by post-mitotic expression of the conserved gene *pdm3*^56^. Further studies focusing on more neural cell types innervating distinct layers and functioning in discrete neural circuits will be essential to relate time and temporal factors expressed in Type II NSCs with the assembly of the circuits in distinct FB layers.

### Hormonal regulation of the neural fate

Our study demonstrates ecdysone signaling via E93 specifies an essential cell type of the sleep-wake circuit, thereby establishing the role of developmental hormonal signaling in specifying sleep-promoting neurons and sleep behavior. These are interesting findings that relate an extrinsic hormonal signal to the stem-cell-intrinsic factors specifying neural fate. Ecdysone signaling plays many roles during development, metamorphosis, and post-development in regulating various physiological processes^110,111,136,137^. Although ecdysone is present at varying concentrations throughout development, cells respond to the ecdysone signaling differentially by expressing unique EcR isoforms temporally. Our recent work demonstrated temporal expression of EcR in Type II NSCs around 55h ALH mediates early to late transition in mid-larval stages^15^. While most animals show the complete loss of dFB neurons, in some animals, we observed a few surviving neurons that are defective in morphology and mistarget to the ectopic layers of the FB (Figure 3F-H’). These data show that EcR acts in Type II NSCs to regulate dFB fate via E93 and might also play essential roles in axonal targeting to the proper FB layers. The FB layers express unique ligand combinations that regulate proper targeting^138^; one likely possibility is that EcR regulates the layer-specific expression of cell adhesion molecules or the receptors for the cell adhesion molecules within the neurons. More studies are needed to show the mechanism of how EcR and ecdysone signaling regulates CX development.

How does E93 regulate cell fate? Ecdysone-induced protein E93 regulates various developmental processes, from wing disc growth to NSC apoptosis^110,119,139–142^. In the developing wing discs, E93 acts as a chromatin remodeler and activates or represses genes by opening and closing chromatin^141^. Further work on identifying E93 targets will help identify the mechanism of E93 function in Type II NSCs. The dFB number did not increase upon the misexpression of E93 in young NSCs, indicating that E93 is not sufficient to specify dFB neurons from the young NSCs. Possibly, a combination of TFs might be required to confer dFB fate. Interestingly, in our misexpression experiments, other Type II lineages did not generate dFB neurons, indicating the role of an unknown spatial factor specific to DL1 NSCs. It is possible that ectopically expressed E93 cannot function in the young NSCs and can only act in the later periods after EcR expression. Previous studies^141^ in *Drosophila* wing discs have shown that not all genomic targets respond to a precocious expression of E93 at all stages, suggesting that temporal factors might be required. We speculate that temporal hormonal signaling might change the chromatin landscape of the late Type II NSCs—as a result, the early/late factors have differential access to the chromatin and thus specify fate by different genes. It will be interesting to investigate the targets of the EcR and E93 in the Type II NSCs and the dFB neurons. In the post-mitotic neurons, E93 might be regulating a battery of effector genes required for consolidating and maintaining dFB neuronal identity, similar to the terminal selector genes^143–145^. The terminal selector genes regulate the expression of effector genes, and an overlap of unique effector genes endows neurons with distinct anatomical and functional properties. Ecdysone and E93 regulate various developmental processes in insects^110,119,142,146^; it will be intriguing to investigate whether E93 regulates the development of tangential input neurons in other insect species.

### Hormonal regulation of sleep behavior

How do developmental hormonal cues specify sleep behavior? Previous studies have reported the role of ecdysone signaling in regulating adult sleep behavior^147–149^. Whole animal ecdysone and EcR hypomorph mutants show adult sleep pattern defects^147,148^. Recently, EcR expression in the cortex glia was also shown to be important in regulating sleep architecture^149^. Our data point to an essential role of developmental hormonal signaling in adult sleep in a cell-type-specific manner via generation of 23E10+ sleep neurons. While evidence over the past decade supports a role for these cells in sleep^50–53,58^, more recent studies suggest that inhibition of these cells has no impact on daily sleep duration in mature adulthood^55,92^ and raise the possibility that the sleep-promoting effect of 23E10+ neuronal activation resides in cells outside the brain^91,92^. Our developmental manipulations contribute further to the apparent complexity of the CX in sleep regulation. E93 knockdown in Type II NSCs causes gross abnormalities in dFB development and morphology while sparing 23E10+ VNC neurons. While mature adult sleep duration is largely intact, sleep consolidation is disturbed, multiple measures of juvenile sleep are impaired, and sleep rebound is abnormal (less consolidated). We propose that the dFB does indeed contain sleep relevant neurons, but not necessarily with a general role in daily sleep control during mature adulthood. Rather, this sleep region of the brain appears to be relevant in conditions when the drive to sleep is heightened, such as in early life or after sleep deprivation. These findings underscore how detailed insights into sleep circuit lineages and development can inform adult sleep regulatory mechanisms.

## Acknowledgments

The authors thank Chris Doe, Doe lab, and Neural Diversity Lab members for their valuable feedback and discussions. Chris Doe and Asif Bakshi for providing critical feedback on the manuscript. We are grateful to Tzumin Lee, Chris Doe, Claude Desplan, Cheng-Yu Lee, and Gerry Rubin for sharing the reagents. Stocks obtained from the Bloomington Drosophila Stock Center (NIH P40OD018537) were used in this study. The monoclonal nc82 antibody was obtained from the Developmental Studies Hybridoma Bank, created by the NICHD of the NIH and maintained at The University of Iowa, Department of Biology, Iowa City, IA 52242. We thank UNM Biology cell biology core for providing the confocal microscopy facility. Company of Biologists Travelling Fellowship to A.R.W, Arnold O. Beckman Postdoctoral Fellowship to J.I-B. The research was supported by NIH DP2NS111996, NIH R01NS120979 to M.S.K and through National Science Foundation CAREER Award IOS-2047020 and Sloan Research Fellowship to M.H.S.

## Author contributions

Conceptualization, A.R.W., M.S.K., and M.H.S.; methodology, A. R. W performed all genetics, anatomy experiments, and confocal imaging with help from G.M.C. K.P performed initial sleep experiments; B. C and J.L performed sleep behavior experiments under the supervision of O.S. and M.S.K.; J. I-B provided the FLPStop2.0 construct and helped design the strategy to generate EcR-FLPStop2.0 flies.; visualization, A.W, M.S.K., and M.H.S.; writing-original draft, A.W, M.H.S; editing, A.W, B.C, M.S.K., and M.H.S.; funding acquisition, M.S.K., and M.H.S.; and supervision, M.S.K., and M.H.S.

## Declaration of interests

The authors declare no competing interests.

## Methods

### Experimental Model and Subject Details

All flies (*Drosophila* melanogaster) were maintained on conventional Bloomington Food formulation at ∼25°C, ∼65% relative humidity, and under a 12-hour light/dark cycle (lights on at 7 AM) throughout development and adulthood unless otherwise stated. The egg-laying for RNAi and FLPStop experiments was performed at 25°C, and then hatched larvae were transferred and allowed to develop at 29°C until adult stages. Also, standardized age matching conversions were used for Gal80^ts^ experiments: 18°C is 2.25X slower than 25°C, and 29°C is 1.03X faster than 25°C^150^. The genotype information of the flies used in each experiment is listed in the Key resource table.

For knocking down E93 in NSCs, we initially used E93 RNAi (from the VDRC stock center) along with UAS Dicer, and the phenotype we observed was severe. Later, we used the same E93 RNAi without dicer, and a similar phenotype was observed. We used a second E93 RNAi from the BDSC stock center, which also gave a similar phenotype.

### Method Details

#### Immunostaining

Fly brains from L3 larvae or 5–7-day old adult flies were dissected in ice-cold insect media (Schneiders media) (Sigma Aldrich) and fixed in 4% paraformaldehyde (PFA) (EMS) in PBST (1X phosphate-buffered saline with 0.5 % Triton X-100) for 27 min at room temperature. Following fixing, three 20-minute washes in 1X PBST were performed, and brains were blocked for 40 minutes in blocking solution PBST containing 2.5% Normal Goat serum and 2.5% Normal Donkey Serum (Jackson ImmunoResearch) at room temperature. After blocking, brain samples were incubated with primary antibody at 4^0^C overnight for larvae and two nights for adult brains. Brains were rinsed and washed thrice for 20 minutes in PBST and then incubated with secondary antibody for 2 hours at room temperature for larvae brains and for adult brains for two nights 4^0^C. After the secondary antibody, brains were rinsed again, and three 20-min PBST washes were performed. After antibody staining, DPX mounting was performed on brain samples. For DPX mounting, the protocol from Janelia FlyLight was followed.

The dilutions for various primary antibodies are as Chicken anti-GFP (1:1500), Rat anti-Dpn (1:500), Rabbit anti-Asense (1:500), Mouse anti-Bruchpilot (nc82) (1:50), Mouse anti-EcR-B1 (1:2000), Rabbit anti-mCherry (1:500), Guinea Pig anti-E93 (1:300).

#### Confocal imaging, data acquisition, and image analysis

Fluorescent image stacks of whole-mount fly brains were taken using a Zeiss LSM 780 confocal microscope. In the figures, only slices corresponding to the FB neuropil (stained with nc82) were shown, giving an exact idea about the axonal targeting of dFB neurons in CX. For dFB neurons stained with GFP, the slices that give the full projection of these neurons were shown. Fiji cell counter plug-in performed adult brain cell counting, and statistical analysis (Student’s T test, one-way ANOVA) was done in Graph Pad prism. Figures were processed and assembled in Adobe Photoshop and Adobe Illustrator, respectively. Asterisks indicate levels of significant differences (*: p<0.05, **: p<0.01, ***: p<0.001, ****: p<0.0001).

#### Clonal Analysis and Birthdating

The CLIn fly females and cell class-specific males were allowed to mate in a bottle and then shifted to an egg-laying cage. The egg laying was done on apple agar caps. The eggs were allowed to hatch at 25°C. Then newly hatched larvae 0-3.5hr old were manually collected and reared on food caps (at 25°C) until the desired time point. For lineage mapping of cell class-specific neurons, larvae were heat shocked at 37°C at Zero ALH. The zero ALH heat shock (for lineage determination) duration was determined and customized to get only one NSC clone one time. The most effective heat shock time was 10-12 minutes at 37 °C to get individual NSC clones. The individual NSC clones obtained were compared to an already available source^64^. For temporal birth dating the neurons from a particular cell class lineage, the 50-minute heat shock was performed at 0h, 48h, and 76h ALH. The larvae were transferred to undergo normal development at 25°C and dissected as adults. Both male and female adult flies were dissected in our experiments.

#### Generation of FlpStop2.0 plasmid for transgenesis

The original FlpStop1.0 was generated to allow for conditional gene control in *Drosophila*^124^. For some genes, for unknown reasons, the FlpStop1.0 cassette did not abrogate expression of the transcript in the presence of FLP Recombinase, though the authors speculated about potential readthrough of transcriptional and translational stop sequences^124^. The FlpStop2.0 cassette was updated to include a 10x repeated sequence of the self-cleaving ribozyme from the Hepatitis Delta Virus (HDV)^151^ (Figure 3A). In other molecular constructs, this sequence has been shown to effectively disrupt transcription and expression of transgenes^125^.

The pFlpStop2.0-attB-UAS-2.1-tdTom plasmid was generated through the synthesis of the FlpStop 2.0 cassette (Figure 3A) and molecular cloning into the FlpStop-attB-UAS-2.1-tdTom^124^ by Genscript (Piscataway, NJ, USA). Constructs were sequence-verified by single primer extension (Sequetech; Mountain View, CA) and were submitted to Addgene. Transgenic flies harboring the FlpStop 2.0 cassette generated through injection of the plasmid and insertion within an intron of the Ecdysone Receptor, as described below.

#### Generation of FlpStop2.0 transgenic flies

Transgenic flies harboring the FlpStop 2.0 cassette in an intron of the Ecdysone Receptor (EcR) were generated via standard construct injection (∼50ng plasmid) by Bestgene (Chino Hills, CA, USA). The MiMIC strain^152^ used for injection was y[1] w[*]; Mi{y[+mDint2]=MIC}EcR[MI05320] (Bloomington Drosophila Stock Center Stock 38619). 4 transgenic lines were identified through the loss of *y* and were PCR-tested for orientation of insertion by Bestgene. We then tested these lines for expression of TdTomato and gene disruption after conditional expression of FLP Recombinase (Figure S3A-D). The transgenic line for conditional disruption of EcR was isolated and has been maintained.

#### Sleep assays

Sleep behavior was measured using the Drosophila Activity Monitoring (DAM) system (Trikinetics, Waltham MA). Single-beam DAM2 monitors were used for sleep deprivation experiments, and multi-beam DAM5H monitors were used for all other sleep experiments. Newly eclosed male flies from crosses were collected and aged in group housing on standard food. Flies were then anesthetized on CO2 pads and loaded into glass tubes (70mm × 5mm × 3mm) containing 5% sucrose and 2% agar medium. Juvenile adult flies were loaded at ∼ZT6 on the day of eclosion; mature adult flies were loaded at the same time but after aging for 6-8 days. To deprive flies of sleep, mechanical shaking stimulus was applied by attaching DAM2 monitors to microplate adapters on vortexers (VWR). Monitors were shaken for 2 seconds randomly within every 20-second window for 12 hours during ZT12 – ZT24. Activity counts were collected every minute, and periods of inactivity lasting at least 5 minutes were classified as sleep; for long sleep bout analysis, only inactivity lasting at least 60 minutes was counted. Sleep parameters were analyzed with a custom R script using Rethomics package^153^.

#### Key Resource Table

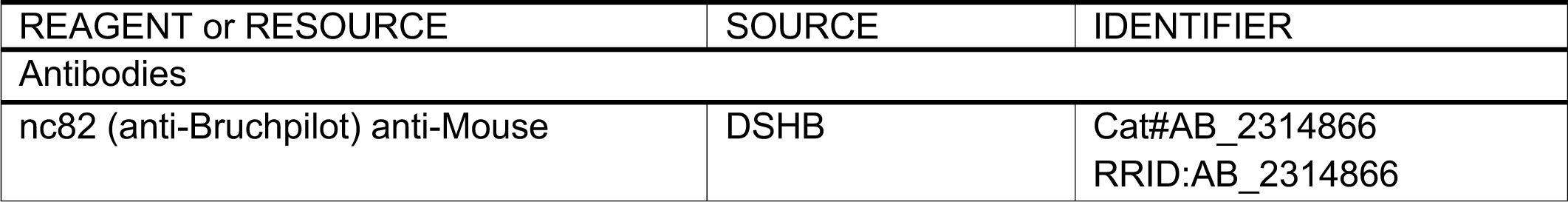
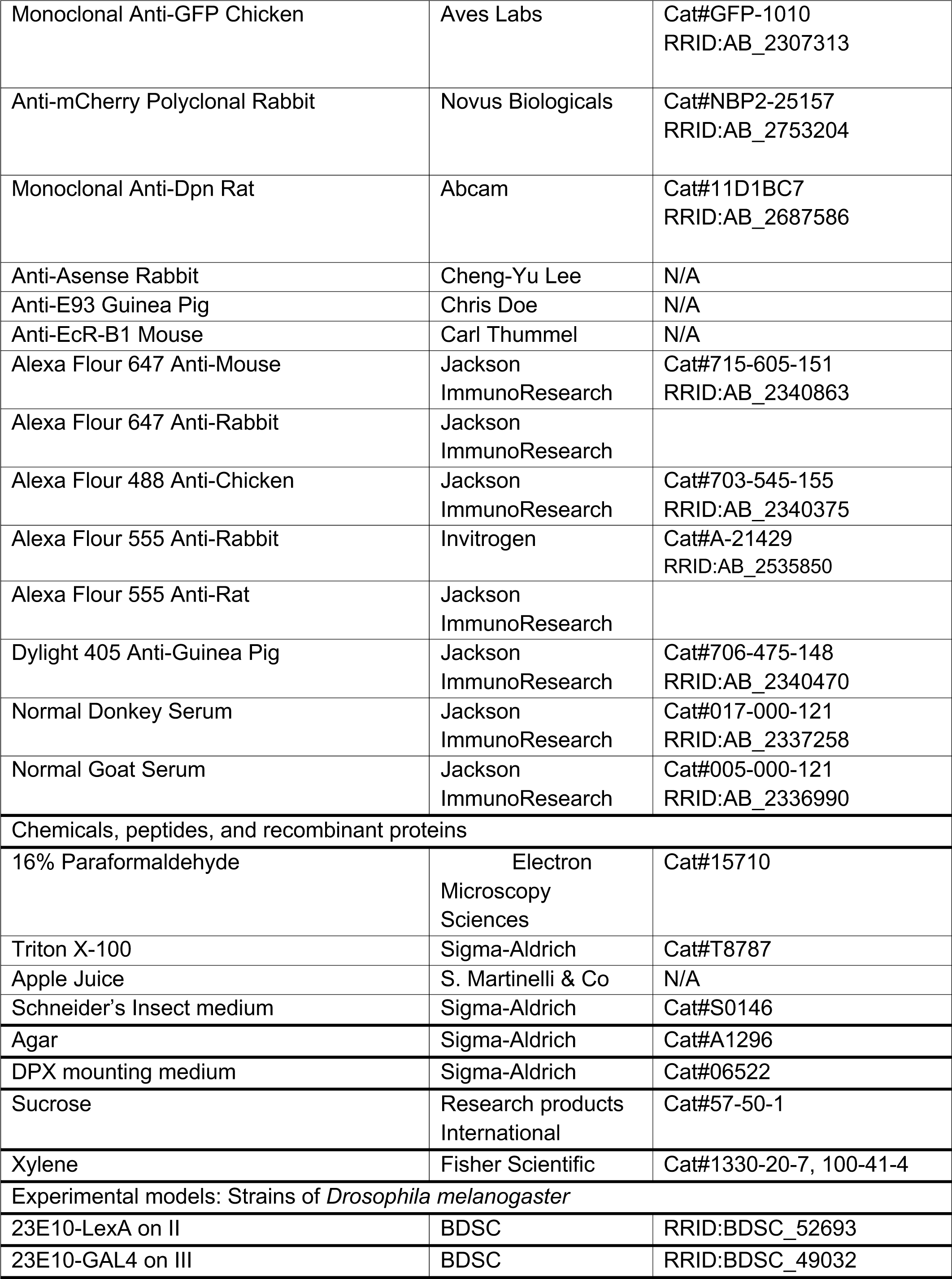
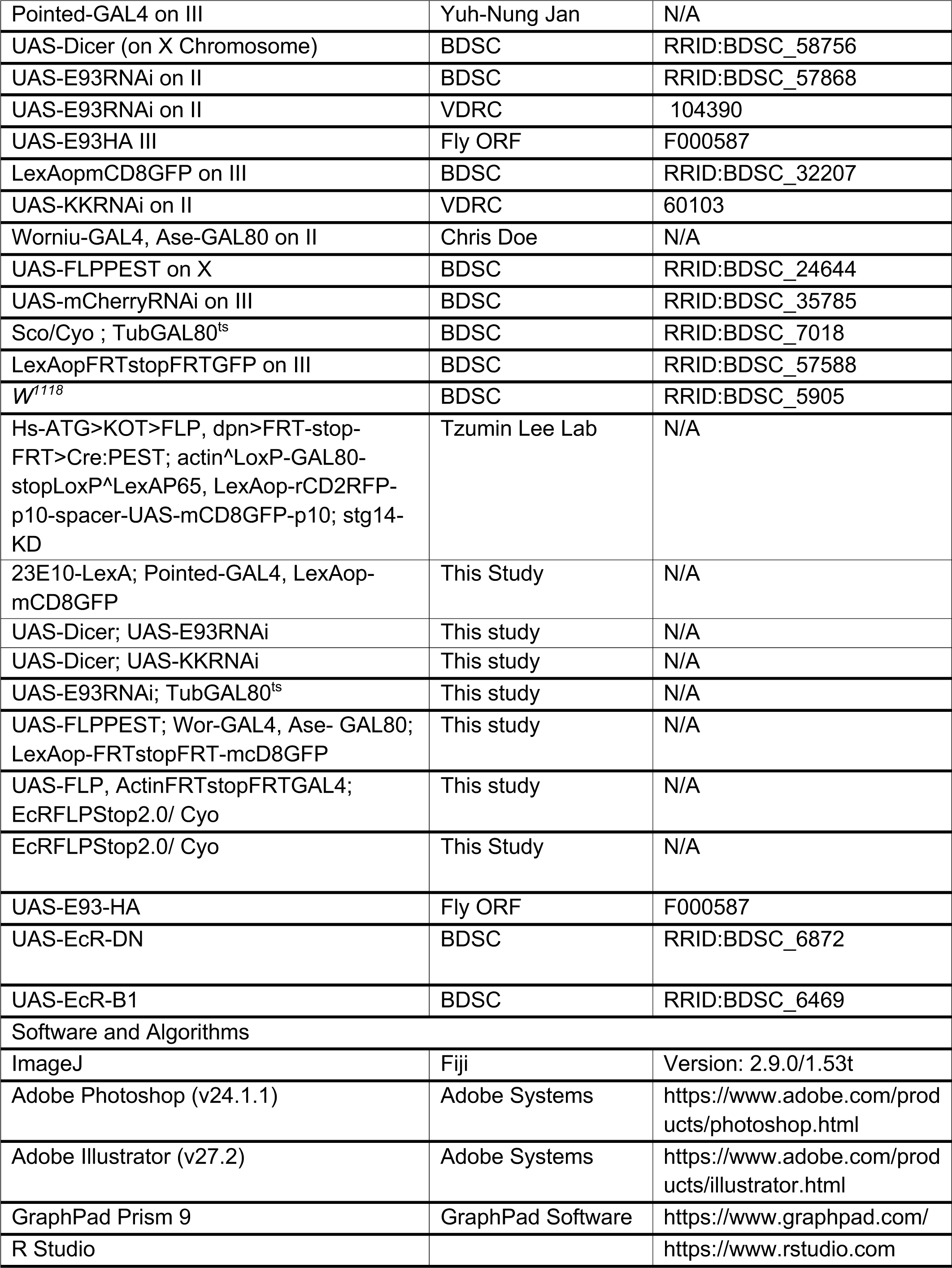
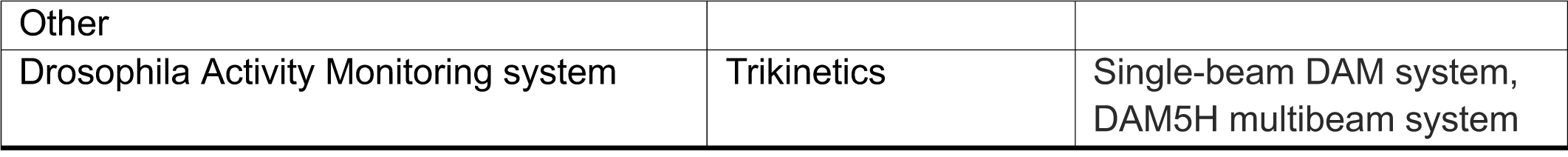

#### Fly Genotypes with Associated Figures

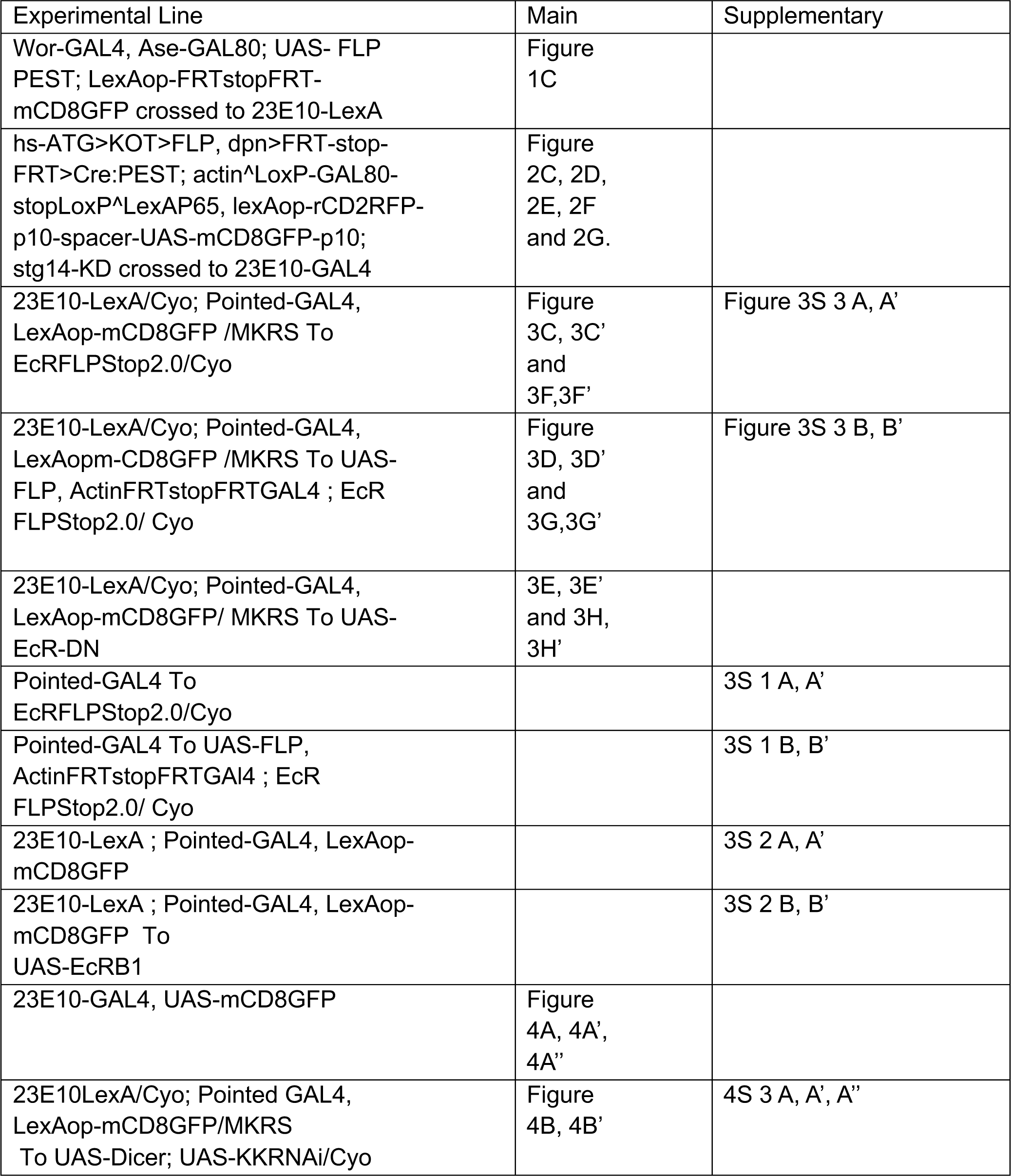
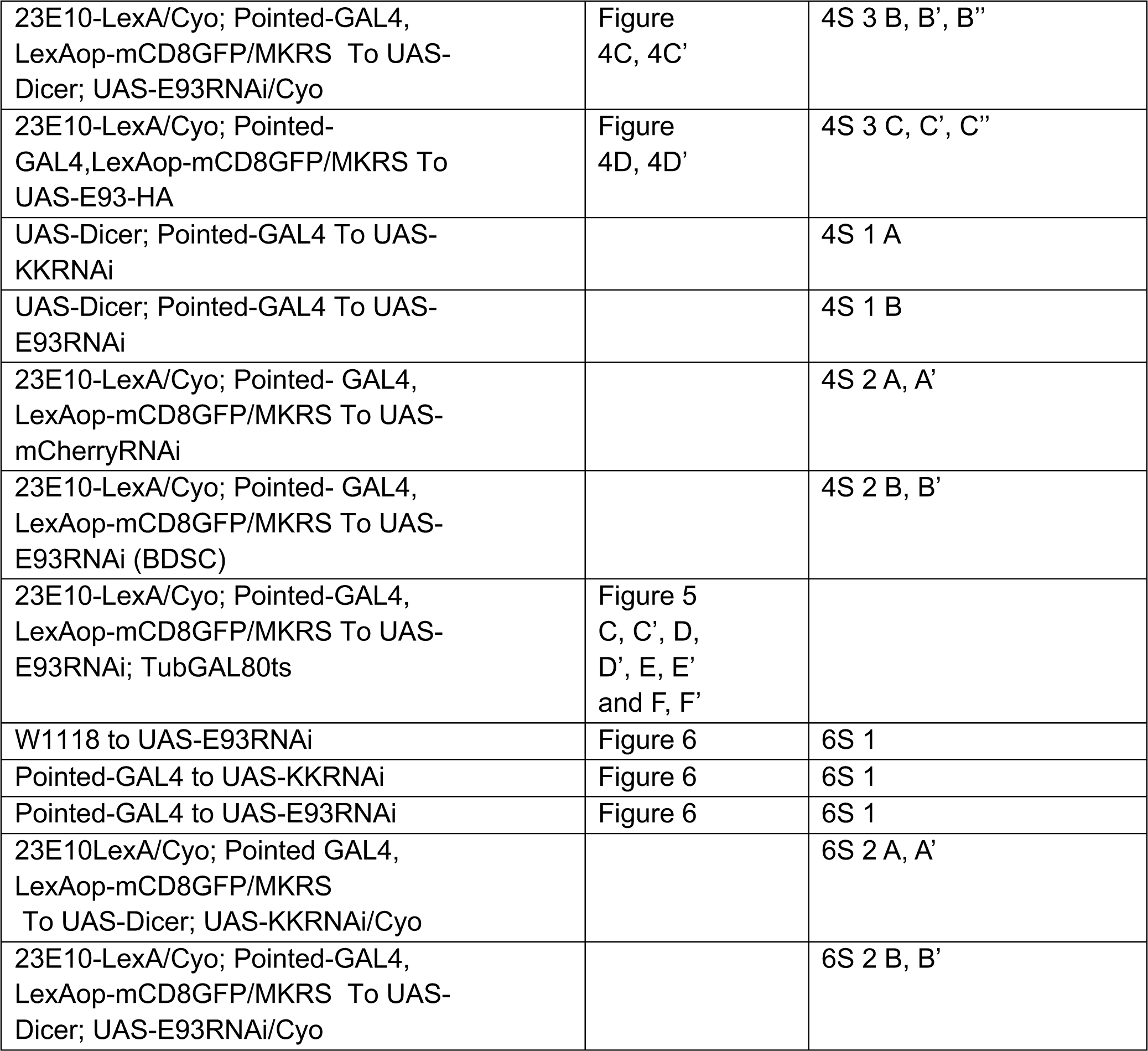

| Experimental Line | Main | Supplementary |
| --- | --- | --- |
| Wor-GAL4, Ase-GAL80; UAS- FLP<br>PEST; LexAop-FRTstopFRT-mCD8GFP crossed to 23E10-LexA | Figure<br>1C |  |
| hs-ATG>KOT>FLP, dpn>FRT-stop-FRT>Cre:PEST; actin<sup>Δ</sup>LoxP-GAL80-stopLoxP<sup>Δ</sup>LexAP65, lexAop-rCD2RFP-p10-spacer-UAS-mCD8GFP-p10;<br>stg14-KD crossed to 23E10-GAL4 | Figure<br>2C, 2D,<br>2E, 2F<br>and 2G. |  |
| 23E10-LexA/Cyo; Pointed-GAL4,<br>LexAop-mCD8GFP /MKRS To<br>EcRFLPStop2.0/Cyo | Figure<br>3C, 3C'<br>and<br>3F,3F' | Figure 3S 3 A, A' |
| 23E10-LexA/Cyo; Pointed-GAL4,<br>LexAopm-CD8GFP /MKRS To UAS-FLP, ActinFRTstopFRTGAL4 ; EcR<br>FLPStop2.0/ Cyo | Figure<br>3D, 3D'<br>and<br>3G,3G' | Figure 3S 3 B, B' |
| 23E10-LexA/Cyo; Pointed-GAL4,<br>LexAop-mCD8GFP/ MKRS To UAS-EcR-DN | 3E, 3E'<br>and 3H,<br>3H' |  |
| Pointed-GAL4 To<br>EcRFLPStop2.0/Cyo |  | 3S 1 A, A' |
| Pointed-GAL4 To UAS-FLP,<br>ActinFRTstopFRTGAI4 ; EcR<br>FLPStop2.0/ Cyo |  | 3S 1 B, B' |
| 23E10-LexA ; Pointed-GAL4, LexAop-mCD8GFP |  | 3S 2 A, A' |
| 23E10-LexA ; Pointed-GAL4, LexAop-mCD8GFP To<br>UAS-EcRB1 |  | 3S 2 B, B' |
| 23E10-GAL4, UAS-mCD8GFP | Figure<br>4A, 4A',<br>4A'' |  |
| 23E10LexA/Cyo; Pointed GAL4,<br>LexAop-mCD8GFP/MKRS<br>To UAS-Dicer; UAS-KKRNAi/Cyo | Figure<br>4B, 4B' | 4S 3 A, A', A'' |
| 23E10-LexA/Cyo; Pointed-GAL4, LexAop-mCD8GFP/MKRS To UAS-Dicer; UAS-E93RNAi/Cyo | Figure 4C, 4C' | 4S 3 B, B', B'' |
| 23E10-LexA/Cyo; Pointed-GAL4, LexAop-mCD8GFP/MKRS To UAS-E93-HA | Figure 4D, 4D' | 4S 3 C, C', C'' |
| UAS-Dicer; Pointed-GAL4 To UAS-KKRNAi |  | 4S 1 A |
| UAS-Dicer; Pointed-GAL4 To UAS-E93RNAi |  | 4S 1 B |
| 23E10-LexA/Cyo; Pointed- GAL4, LexAop-mCD8GFP/MKRS To UAS-mCherryRNAi |  | 4S 2 A, A' |
| 23E10-LexA/Cyo; Pointed- GAL4, LexAop-mCD8GFP/MKRS To UAS-E93RNAi (BDSC) |  | 4S 2 B, B' |
| 23E10-LexA/Cyo; Pointed-GAL4, LexAop-mCD8GFP/MKRS To UAS-E93RNAi; TubGAL80ts | Figure 5 C, C', D, D', E, E' and F, F' |  |
| W1118 to UAS-E93RNAi | Figure 6 | 6S 1 |
| Pointed-GAL4 to UAS-KKRNAi | Figure 6 | 6S 1 |
| Pointed-GAL4 to UAS-E93RNAi | Figure 6 | 6S 1 |
| 23E10LexA/Cyo; Pointed GAL4, LexAop-mCD8GFP/MKRS To UAS-Dicer; UAS-KKRNAi/Cyo |  | 6S 2 A, A' |
| 23E10-LexA/Cyo; Pointed-GAL4, LexAop-mCD8GFP/MKRS To UAS-Dicer; UAS-E93RNAi/Cyo |  | 6S 2 B, B' |

## Notes

### Competing Interest Statement

The authors have declared no competing interest.

