## Supplemental Figures for "Stem cell-specific ecdysone signaling regulates the development and function of a *Drosophila* sleep homeostat"

Figure 3S 1

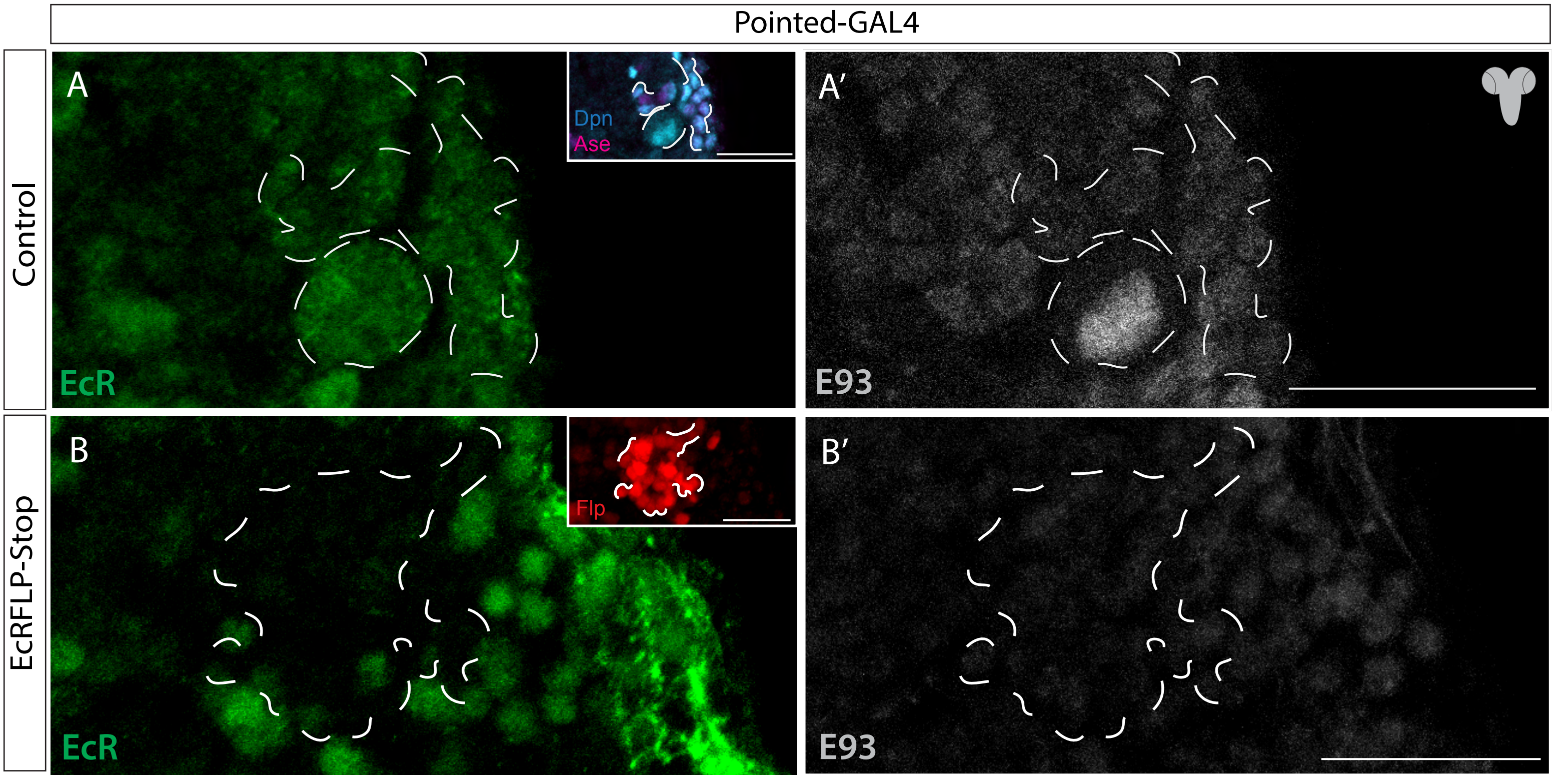


**Figure 3S 1**
**EcR-FLPStop is sufficient to reduce EcR expression in Type II NSCs.**
A-A') EcR and E93 are expressed in L3 larval Type II NSC lineages (outlined). NSCs are identified by Dpn+ Ase- expression (see inset). B-B') EcR-FLPStop triggered by FLP expression in Pointed-GAL4 pattern leads to a loss of EcR and E93 in L3 larval Type II NSCs. The inset (red color) shows the FLP-based disruption happening in the Type II lineages.
Scale bars, 20  m, n = 14 larval brains.

Figure 3S 2

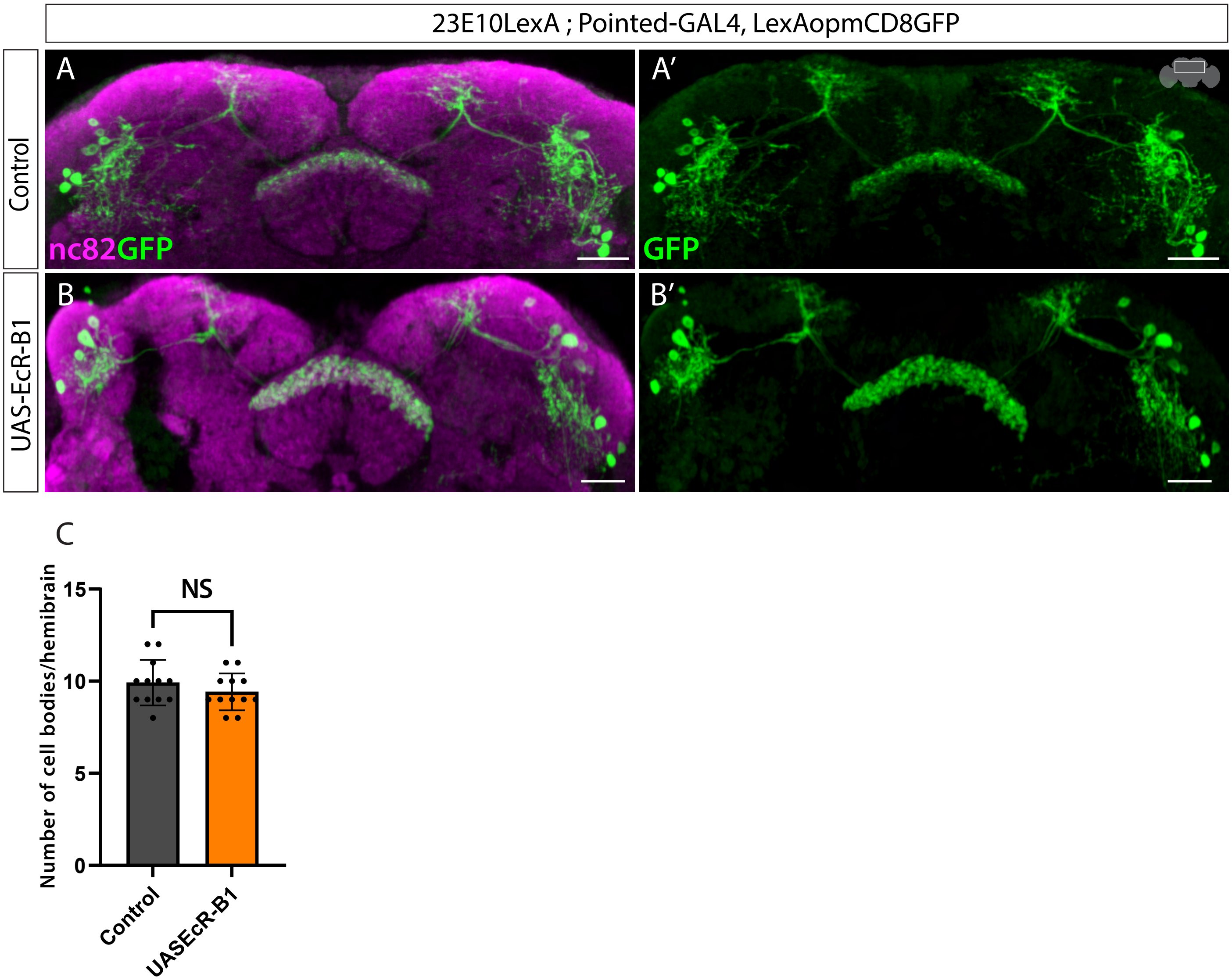


**Figure 3S 2**
**EcR is not sufficient to generate ectopic dFB neurons**
A, A') dFB neurons labeled by a GFP reporter in control brains. B, B') dFB neurons labeled by GFP upon EcR-B1 overexpression in Type II NSCs showing no change in cell body number or morphology. C) Quantification of the number of cell bodies per hemibrain. Error bars represent SEM; * $p<0.05$, ** $p<0.01$, *** $p<0.001$, **** $p<0.0001$, NS, non-significant by Student t-test. Scale bars, 20μm. n=12 adult hemibrains.

Figure 3S 3

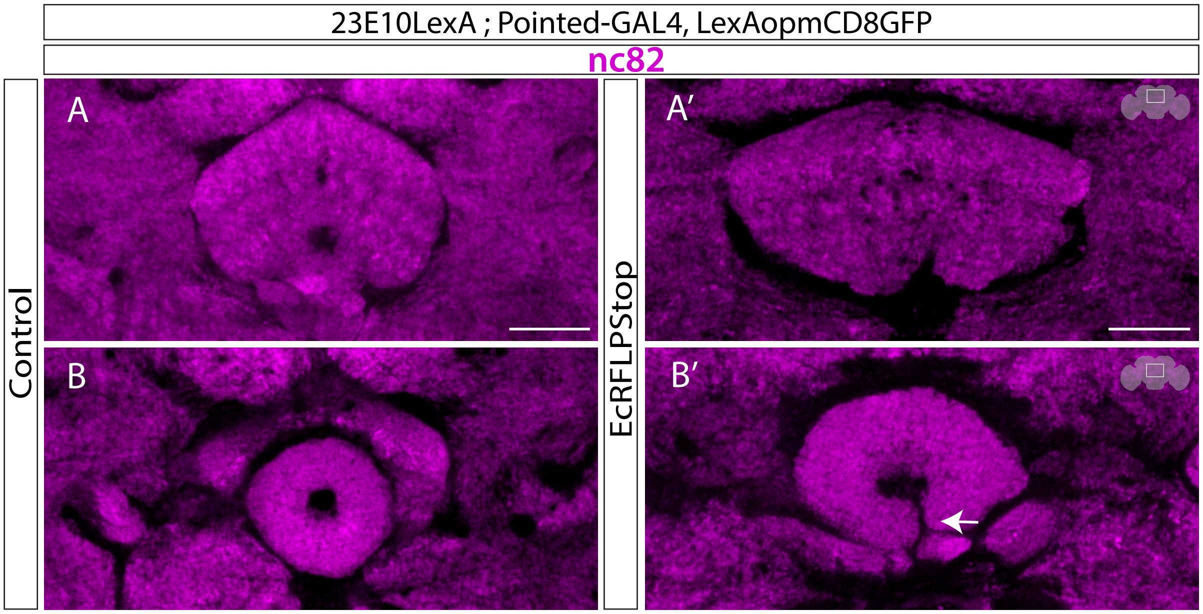


**Figure 3S 3**
**EcR is required for normal EB development.**
A, B) The morphology of the neuropil structures, FB, and EB in control brains.
A’, B’) Type II NSC-specific EcR loss of function results in defective morphology of the EB while the FB is normal.
Scale bars 20μm, n = 6 adult brains. nc-82 labels the neuropil structures.

Figure 4S 1

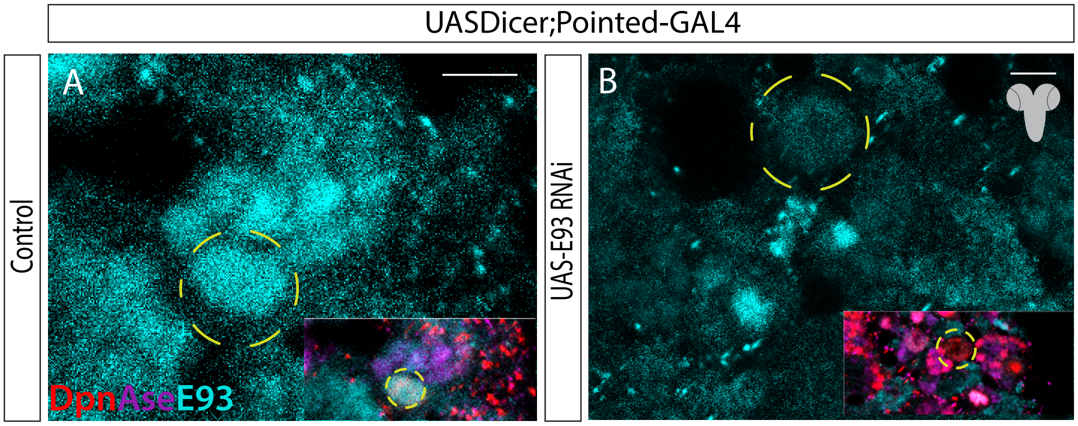


**Figure 4S 1**
**E93 RNAi expression in Type II NSCs knocks down protein expression effectively.**

A) Shows expression of E93 in Type II NSCs.
B) Significant loss of E93 in Type II NSCs upon E93 RNAi expression. Small panels in both brains depict the Dpn and Ase staining, which are used to locate the Type II NSCs. Scale bars, 5µm n= 4 larval brains.

Figure 4S 2

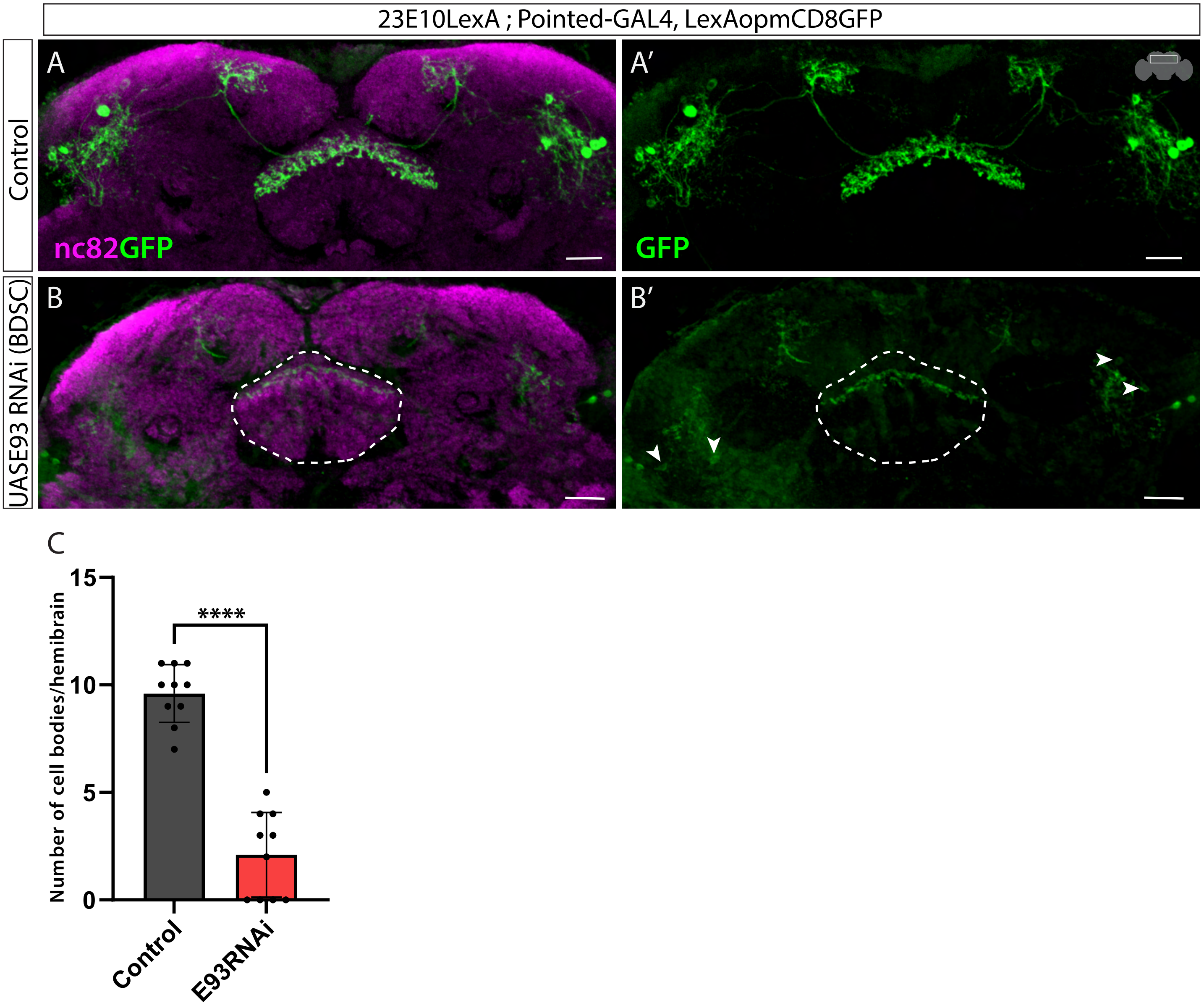


**Figure 4 S2**
**E93 phenotype is specific to E93 knockdown.**
A, B') Compared to control (A, A'), the dFB neurons are not specified in E93RNAi (BDSC) animals. Two E93RNAi lines from two different sources produce similar phenotypes.C) Quantification of the number of cell bodies per hemibrain. Error bars represent SEM; * p<0.05, **p<0.01, ***p<0.001, **** p<0.0001, NS, non-significant by Student t-test. Scale bars, 20μm. n=10 adult hemibrains.

Figure 4S 3

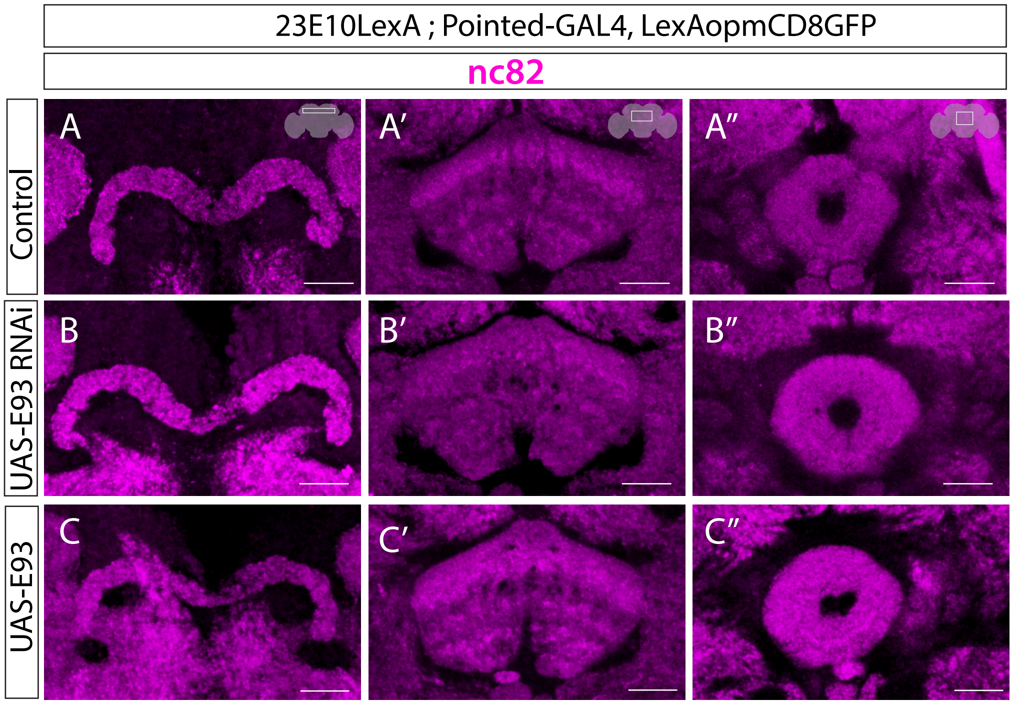


**Figure 4 S3**
**E93 in Type II NSCs is not necessary for the CX neuropil development**
A-B’’) The morphology of three CX neuropil structures, PB, FB, and EB, in control animals (A-A”) is not significantly different in E93RNAi animals (B-B’’). C-C”) shows PB, FB, and EB neuropils in E93 over-expression brains. Scale bars, 20  m, n = 5 adult brains.

Figure 6S 1

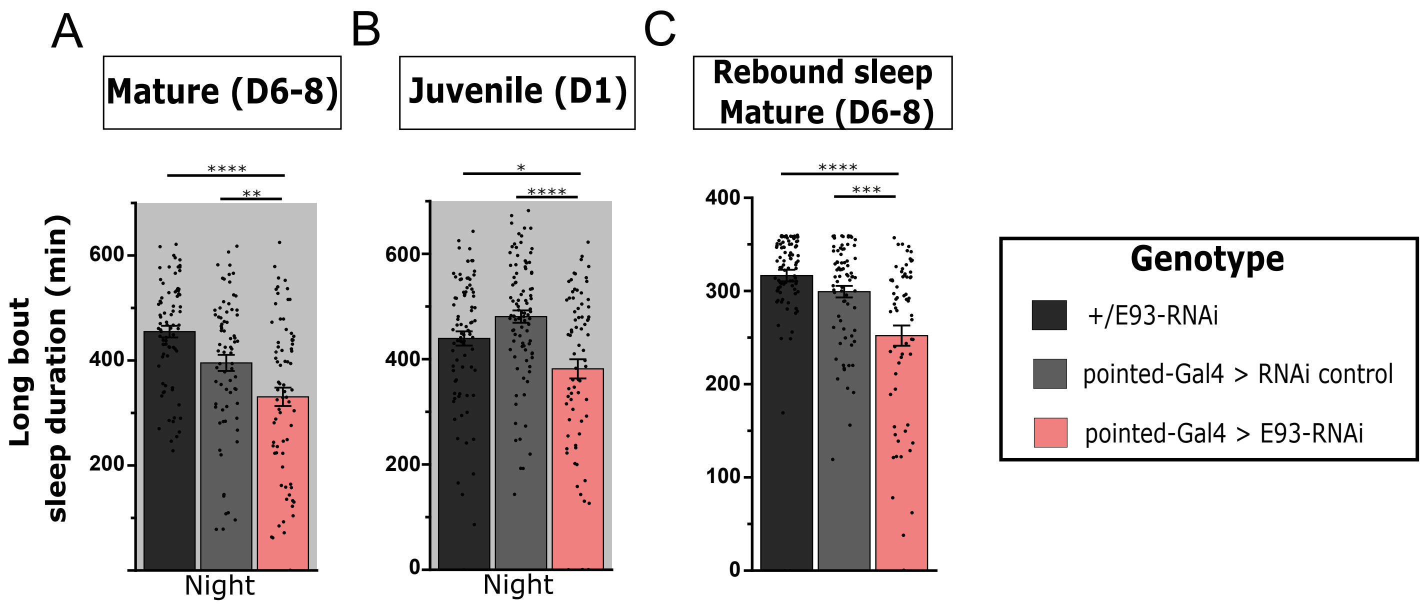


**Figure 6 S1**

**E93 knockdown in Type II NSCs reduces sleep comprised of long bouts in adulthood**

Quantification of sleep duration comprised of long sleep bouts (bouts >60 minutes) during the night in (A) mature adults and (B) juvenile adults or (C) during the day (6 hours) following a night of sleep deprivation in mature adults. In each case, the genotype is E93-RNAi under the control of Pointed-GAL4 (red) compared to genetic controls (black, gray). n=79,74,74 left to right (A), n=85,94,74 (B), and n=93,82,67 (C). Error bars represent SEM; *p<0.05, **p<0.01, ***p<0.001, ****p<0.0001 by One-way ANOVA with Mann-Whitney multiple comparisons test corrections using Holm method.

Figure 6S 2

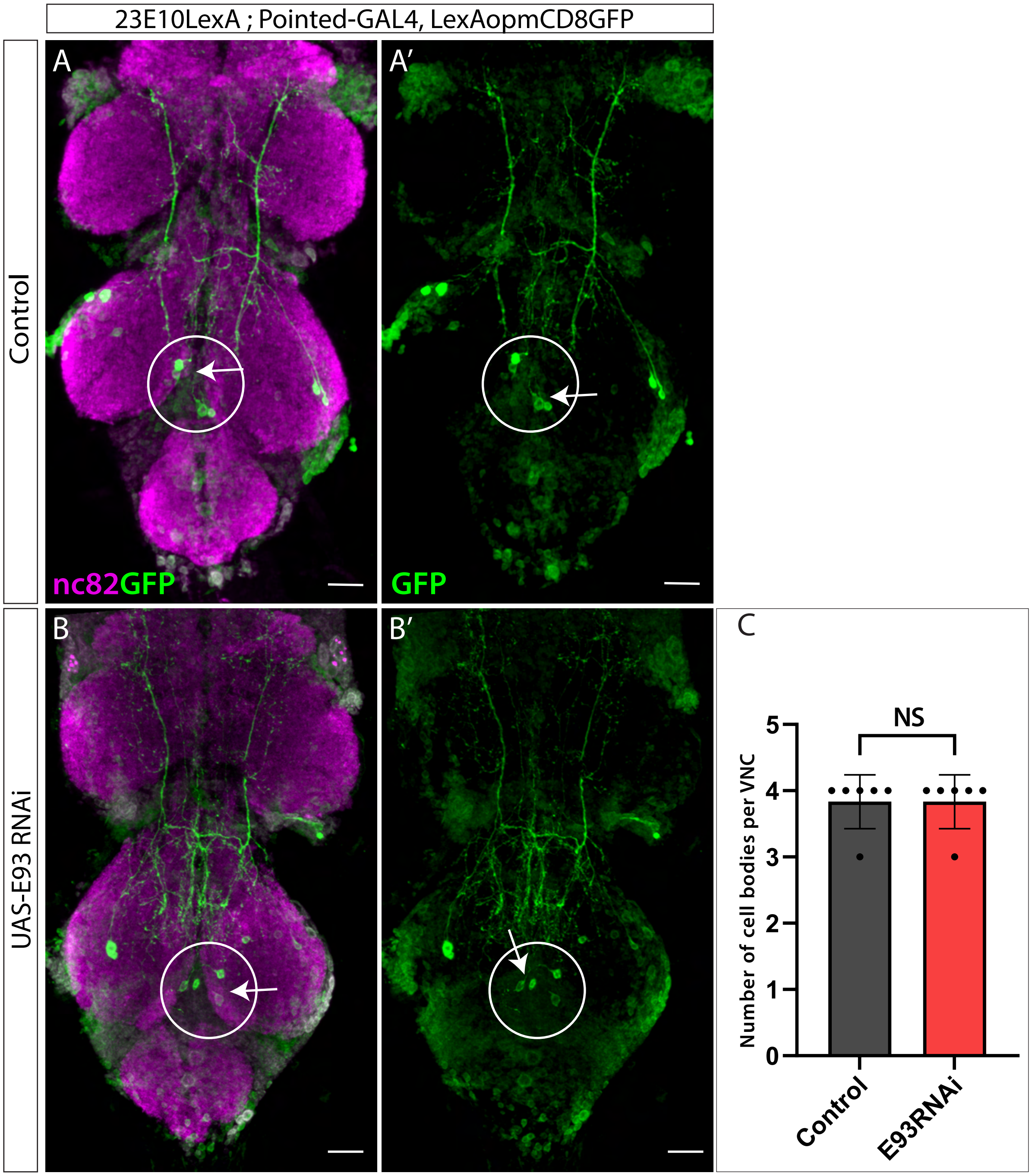


**Figure 6 S2**

**E93 RNAi knock-down in Type II NSCs does not affect 23E10+ neurons located in adult VNC**

A, B') shows cell bodies of VNC neurons in control flies.

B, B) no change in the VNC cell body numbers upon E93 knockdown.

C) Quantification of the number of cell bodies per VNC in control and RNAi flies. Error bars represent SEM; * p<0.05, **p<0.01, ***p<0.001, **** p<0.0001, NS, non-significant by Students t-test.

Scale bars, 20µm, n = 6 adult brain VNCs.
